# YAP1 Contributes to The Development of Contractile Force and Sarcomere Maturation in Human Pluripotent Stem Cell-Derived Cardiomyocytes

**DOI:** 10.1101/2024.07.02.601803

**Authors:** Vladimir Vinarsky, Stefania Pagliari, Fabiana Martino, Cristina Mazzotti, Katerina Jirakova, Zuzana Garlikova, Enrico Di Iuri, Daniel Kytyr, Patrizia Benzoni, Martina Arici, Alessia Metallo, Kira Zeevaert, Andrea Barbuti, Wolfgang Wagner, Marcella Rocchetti, Giancarlo Forte

## Abstract

**Background:** Perspective regenerative therapies for injured myocardium focus on reactivating developmental processes to regenerate damaged contractile tissue. In animal models, the Hippo pathway was shown to improve heart regeneration after myocardial infarction, possibly by expanding the pool of cardiomyocytes. We hypothesized that activating the Hippo pathway’s downstream effector, Yes Associated Protein (YAP1), may have effects beyond promoting proliferation in human cardiomyocytes. We have therefore investigated how YAP1 influences human cardiomyocyte maturation, sarcomere structure, electrophysiological properties, their response to mechanical stimuli, intracellular Ca^2+^″ dynamics and force development using models of cardiomyocytes derived from pluripotent stem cells.

**Methods:** We employed genetic models of YAP1 deficient human embryonic and induced pluripotent stem cells, cardiomyocyte differentiation, compliant cultivation substrates, mechanical actuation, ECM deposition, super resolution microscopy, electrophysiological measurements and engineered heart tissues (EHTs) to characterize the effects of YAP1 deficiency on cardiomyocytes during maturation. We also used full length YAP1 re-expression to rescue the effects of YAP1 deficiency in contracting cardiomyocytes.

**Results:** YAP1 contributes to cardiomyocyte maturation, participates in the formation and alignment of myofibrils, as well as in the maturation of electrophysiological properties. The net effect of YAP1 deficiency in cardiomyocytes is the inability to respond to physiological stimuli by compensatory growth resulting in reduced force development. Additionally, YAP1 reactivation in contracting cardiomyocytes leads to rescue of myofibril maturation.

**Conclusions:** This research demonstrates that YAP1 activity is essential to promote cardiomyocyte maturation, contractility, and response to regeneration inducing stimuli.

## Introduction

The reduced contractile activity of the diseased heart is largely ascribed to the loss of functional cardiomyocytes (CMs). While lower vertebrates like zebrafish and newts reveal adult heart regeneration even after extensive injury, damages to the adult human heart only promote a reparative response ^1^. This is largely due to the terminally differentiated state of the human post-natal cardiomyocytes, which are not able to proliferate and to the lack of endogenous stem cells ^2^. While the implantation of *in vitro* generated cardiomyocytes has gained momentum after the discovery of induced pluripotent stem cells ^3^, the stimulation of the endogenous proliferative capacities of cardiomyocytes is at present among the most promising avenues to reach the bedside. Reactivating the proliferation of resident cardiomyocytes or increasing their contractility hold promise to restore heart functionality - by well-integrated cardiomyocytes attuned to their environmental cues. To this end the Hippo pathway and its effector Yes Associated Protein 1 (YAP1) have recently become prime targets of investigations due to their role during development and adult heart function ^4^.

Together with its paralog protein WWTR1, YAP1 sustained nuclear presence is considered largely responsible for the initiation, growth and dissemination of most solid tumors ^5^. These effects are obtained by directly or indirectly targeting the expression of genes involved in cell differentiation ^6^, proliferation^7^, and migration^8^. The nuclear exclusion and degradation of this ubiquitous transcriptional co-factor is mostly regulated by Hippo pathway ^9^. In brief, Hippo member LATS1/2 phosphorylates YAP1 on S127 and several other residues in transactivation domain, which leads to 14-3-3 dependent and independent cytoplasmic retention and degradation ^10^. The nuclear localization of YAP1 leads to its association with transcriptional partners and the activation of gene expression in a context-specific fashion. TEAD family of transcriptional factors was shown to be the major mediator of YAP1 activity ^11^. While several Hippo independent mechanisms that regulate YAP1 localization through F-Actin assembly, Rho, AMPK, and Erbb2 were described ^12, 13^, Hippo-YAP1-TEAD axis is the prevalent and most understood mode of action.

The activity of Hippo-YAP1 pathway during development is multiphasic. Initially, high activity of YAP1 protects pluripotency in stem cells ^14^. Later, to enable mesoderm specification its expression is repressed, to be again re-activated to drive the proliferation of embryonic cardiomyocyte progenitors ^15^. The absence of either YAP1 or TEAD at the progenitor expansion stage leads to hypoplastic heart and embryonic death ^16, 17^. In adult heart, Hippo mediates the nuclear exclusion of YAP1 to stop cardiomyocyte proliferation and prevent cardiomegaly ^18^. Still, finely tuned YAP1 activity is required for adult heart function. For example, while perinatal deletion of TEAD1 ^19^ or YAP1 in cardiomyocytes leads to dilated cardiomyopathy after increased load or injury ^20, 21^, so does constitutive repression of Hippo pathway or TEAD overexpression after pressure overload ^22, 23^. Significant efforts are being made to identify and exploit subsets of YAP1 targets driving cardiomyocyte proliferation since the discovery that YAP1 sits at the crossroad of a handful of microRNAs being able to reactivate adult CM proliferation ^24, 25^.

Cardiac pathologies induce continuous remodeling of ECM characterized by deposition of fibronectin, increased ECM stiffness and strain. Adverse cardiac remodeling drives a major shift in cardiomyocyte contractile function to favor cardiac cell survival and preserve heart pumping function ^26^. Convincing *in vivo* evidence shows that cardiac cells respond to the ensuing changes in the composition and mechanical stress caused by ECM remodeling by YAP1 nuclear re-expression (Del Re 2013; Mosqueira 2014). While proliferation is the main observed effect of YAP1 activity in fetal heart, cell survival and hypertrophy become dominant in the adult heart ^20, 21, 27^. Further *in vitro* work showed the importance of cell-ECM interaction, actin polymerization, and mechanical stimulation on YAP1 dependent hypertrophic response, which is not fully corroborated by data from the overexpression of Hippo-insulated YAP1 S127A variants or Hippo pathway inhibition.

Our group has previously investigated the effects of ECM mechanics on YAP1 activity. We found that rearrangement of ECM 3D structure in human failing heart prompts the re-expression of YAP1 in the nuclei of cardiac fibroblasts and contributes to their proliferation ^28^. In addition, we showed that ECM remodeling fine-tunes YAP1 transcriptional activity in the nucleus of the cardiomyocytes of diseased human heart by its alternative mRNA splicing ^29, 30^. Nonetheless, the effects of YAP1 activity on the transcriptional landscape of terminally differentiated cardiomyocytes are underexplored.

We here took advantage of YAP1-KO human embryonic and induced pluripotent stem cells (hESCs and hiPSC, respectively) to investigate the role of endogenous YAP1 on the expression of major molecular markers, morphology, sarcomere maturation, electrophysiological properties and intracellular Ca^2+^ dynamics. We show that YAP1 deficiency disrupts the development and maturation of CMs contractile and excitation-contraction (EC) coupling apparatus, a phenomenon that can be partially rescued by YAP1 re-expression in beating CMs. In addition, we highlight how YAP1 absence abolishes the compensatory response to biochemical and mechanical stimulation. Finally, we demonstrate that the complex effects of YAP1 deficiency on the excitation and contraction apparatus determine a significant reduction in the force generated by engineered heart tissues (EHTs).

This work shows novel observations of YAP1 role in CM maturation and physiology, which could be leveraged for therapeutic use in heart failure.

## Materials and methods

### Cell Culture

#### Cell-lines differentiation

The YAP deficient (YAP1-KO) and isogenic H9 (WT or CTRL) human embryonic stem cell lines (hESCs) were a kind gift of Miguel Ramalho-Santos and Han Qin ^14^. The cells were maintained in an undifferentiated state by culturing them on Matrigel Growth Factor Reduced (1:100 in DMEM/F12, Corning, NY, USA) coated tissue culture plastic plates in complete Essential 8^TM^ Medium (E8, Thermo Fisher Scientific) containing penicillin/streptomycin (0.5%, VWR).

The YAP deficient iPSC cells and the isogenic wild type (hPSCreg ID: UKAi009-A) have been characterized and described in detail before ^31^. In brief, the iPSCs were derived after informed and written consent using guidelines approved by the Ethic Committee for the Use of Human Subjects at the University of Aachen (permit number: EK128/09). A CRISPR/Cas9 nuclease approach was then used to target exon three of the human YAP1 gene, which is shared in all transcripts. iPSC lines were initially cultured on tissue culture plastic coated with vitronectin (0.5 µg/cm^2^; Stemcell Technologies, Vancouver, Canada) in StemMACS iPS-Brew XF (Miltenyi Biotec GmbH, Bergisch Gladbach, Germany).

Cell differentiation was conducted using previously described protocol ^32^ with slight modifications by using sequential inhibition of GSK-3 by CHIR99021 (8 µM, Sigma-Aldrich) and WNT by IWP-2 (5 µM, Selleck chemicals) in RPMI 1640 media (Gibco) supplemented with B27 supplement without insulin (RPMI +B27 -Ins) (1x, Gibco), containing penicillin/streptomycin (1%, VWR). Geltrex^TM^ (1:100, Gibco) was added to cell culture media during the first six days of the differentiation. When the cell cultures started contracting the RPMI media was supplemented with B27 containing insulin (RPMI +B27 +Ins) (1x, Gibco). The culture media were exchanged every 2-3 days. Cells on differentiation days 10, 15-20 and 30-40 were used for experiments.

#### Single-cell cardiomyocyte culture

At specified timepoints the confluent cell cultures were enzymatically dissociated using Multi Tissue Dissociation Kit 3 (Miltenyi Biotec) and replated at 10.000 cells/cm^2^ in RPMI +B27 +Ins media with ROCK inhibitor Y27632 (5 µM, Selleck chemicals) supplemented RPMI media to facilitate cell attachment. The media was changed the next day for RPMI +B27 + Ins media without ROCK Inhibitor. The experiments on single cell cultures were performed on day four after replating. The fourth day timepoint was selected as a day when the replated single cell cardiomyocytes resumed spontaneous contraction.

#### Surface coating

Single cardiomyocyte cell culture dishes were coated with Matrigel Growth Factor Reduced (1:20 in RPMI, Gibco, Corning). For ECM concentration experiments and mechanical actuation experiments on Mechanoculture FX2 plates fibronectin coating (1 µg/mL and 10 µg/mL, in PBS StemCell Technologies) at 37°C for 1 hour were used.

#### Cell treatments

For actin tension inhibition experiments, single cardiomyocytes were treated with latrunculin A (250 nM, Cayman Chemical) for 24 hours.

#### PDMS-preparation

Mixtures of Sylgard 184 and Sylgard 527 were used to prepare compliant materials with Young modulus 10 kPa as published previously ^33, 34^. The stiffness was tested as described previously ^34^.

#### Mechanical actuation protocol

The mechanical stimulation experiments were conducted on single cell cardiomyocyte cultures seeded on compliant stretching plates supplied with Mechanoculture FX2 device (CellScale). Confluent cultures of beating cardiomyocytes at day 15-20 of differentiation were enzymatically dissociated and seeded at 10.000 cells/cm^2^ on the fibronectin coated stretching plates. After four days of single cell culture, the cell substrate was stretched to 120% of the well length during one second and kept static for next 24 hours, before fixation with 4% PFA followed by immunofluorescence staining.

#### Engineered Heart Tissues (EHT) and measurements of contractile force

EHTs were generated using Cuore device (Optics11-Life, Amsterdam, NL), a pillar-based system for standard 24-well plates that uses integrated fibre optical sensing and Electrical Pulse Stimulation (EPS) for continuous electrical stimulation and real-time recording of contractile activity ^35^. The 24-well casting plate was pretreated O/N with 1% Pluronic acid (Sigma) and kept at 4 °C for 24 hours to prevent cell attachment. The day after, 0.5 × 10^6^ WT and YAP1-KO hiPSC-CMs at day 30 of differentiation were mixed, on ice, with Collagen type I (0,072 mg/EHTs, TeloCol®-6, Advanced Biomatrix), 0,003 N NaOH and RPMI +B27 + Ins supplemented with 20% FBS and 5 µM Y27632. The cell-hydrogel mixture was distributed into 24-well casting plate and placed inside the Cuore device equipped with an optical fibers plate containing an array of 24 couples of cantilevers. The device was then placed inside the incubator in order to allow the condensation of the cell/hydrogel mixture for 1 hour and then more media were added. Twenty-four hours later, the media were substituted with RPMI +B27 + Ins. Three/four days from the casting, the array of cantilevers with the attached EHTs was transferred into a new 24-well plate (Corning™ Costar™). Half of the medium in each well media was replaced every 48h hours. were kept in culture for 10 days. The estimated contractile force generated from WT and YAP-KO EHTs was recorded at different time points from day 6 to day 10. The data were analysed using an integrated computer software to calculate the absolute contractile force and the beating.

#### Viral particles production

Plasmid containing full length YAP1 with T2A-mCherry tag (#74942, Addgene) was transfected together with envelope expressing plasmid 9 - PMS2.G (#12259, Addgene), and empty backbone packaging plasmid PSPAX2 (#12260, Addgene) into HEK293T cells using FuGENE® HD Transfection Reagent according to manufacturer’s instructions. The media with HEK293T produced viral particles was collected daily for 4-5 days post transfection and concentrated using Vivaspin20 100kDa protein concentrators to achieve 30x concentrated stock of viral particles. The YAP1 deficient cardiomyocytes were transduced using 8x concentrated viral particles in presence of Polybrene Transfection Reagent (7,5 µg/mL, MERCK). After visual confirmation of YAP1 expression by means of mCherry at day four post transduction, cardiomyocytes were replated into the single cardiomyocyte cell cultures for experiments.

### Immunofluorescence

#### Adherent cultures

Immunofluorescence staining was executed as published previously^36^. In detail, cells were fixed by 4% PFA for 10 minutes, washed and stored in PBS prior to staining. Permeabilization and blocking of non-specific binding epitopes were done using 0.1% and 0.05% Triton X-100 and Tween-20 respectively in 1% BSA in PBS. Incubation with primary antibodies at 4°C overnight in 0.05% Tween-20 in 1% BSA in PBS was followed by washes in 0.05% Tween-20 PBS before incubation with secondary antibodies labelled with fluorescence labels. For primary and secondary antibodies see Supplementary Table 1.

#### Engineered heart tissues

The samples were fixed in 4% PFA for 1 h at room temperature (RT) and then incubated with 15% sucrose solution (Sigma Aldrich) supplemented with 0.03% eosin (Sigma Aldrich) overnight at 4°C. EHTs were then embedded in OCT solution (Leica), frozen in cassettes embedded in isopentane (VWR) cooled with dry ice and stored at −80℃ until cryosectioning. The frozen EHTs were cryosectioned using the CryoStar NX70 Cryostat (ThermoFisher Scientific). Frozen sections were cut at 10 μm thickness, placed onto Menzel Gläser, SuperFrost® Plus slides (ThermoFisher Scientific) and stored at −20℃ until their immunostaining following the same protocol as described above. The slides were mounted with Fluoromount-G (Invitrogen). For primary and secondary antibodies see Supplementary Table 1.

#### EdU proliferation assay

Single cell cultures of cardiomyocyte were exposed to base analogue EdU (10 µM, BaseClick) in cell culture medium for thirty minutes. Following the exposure, cell cultures were washed with PBS and fixed for immunofluorescence staining. The EdU visualization preceding immunofluorescence staining was performed according to manufacturer instructions.

### Western-Blotting

Confluent cell cultures of spontaneously beating cardiomyocytes were enzymatically dissociated using Tryple enzyme and collected in cold PBS. The pellets were snap frozen and stored at −80°C. The pellets were resuspended in 1% SDS lysis buffer as described previously ^37^. Protein concentration was measured using Pierce™ BCA Protein Assay Kit (Thermofisher Scientific) and normalized to 1 mg/mL. Laemli was added to the samples and they were denaturated at 95°C for five minutes. Twenty micrograms of protein per sample were loaded into each lane of 4-20% Mini-PROTEAN® TGX™ Precast Protein Gels (Bio-Rad). The proteins were transferred to nitrocellulose membrane using semidry Trans- Blot® Turbo™ Transfer system (Bio-Rad) and immunodetected using specific primary antibodies diluted in 5% BSA or 5% low fat milk overnight. The signal of secondary HRP conjugated antibodies was visualized by Clarity Western ECL Substrate (Bio-Rad) and acquired using ChemiDoc XRS+ system (Bio-Rad). Integrated density of the signal of detected protein was quantified using Image Lab Software (Bio-Rad). See Supplementary Table 1 for list of used antibodies.

### Image acquisition

The confocal microscope ZEISS LSM780 (Zeiss, Oberkochen, Germany) was used to acquire images using 10x magnification for tile scans, 40x for cell size measurement and YAP1 localization, and 63X magnification for sarcomere length analysis according to manufacturer’s description. ZEISS Elyra 7 with lattice SIM module was used to acquire images with sub-0.1 µm resolution in all three dimensions. The SIM processing was done using manufacturer algorithm.

### Image analysis

#### Cell area and YAP1 localization

Cell area and YAP1 localization was measured using a combined pipeline of Ilastik ^38^ and CellProfiler 4 ^39^ custom pipeline. The cardiac troponin channel was used to segment cardiomyocytes (cell object), DAPI channel to segment nuclei. The nuclei objects were subtracted from the cell objects to create a cytoplasm object. For all objects area, circularity, and intensity of immunofluorescence of YAP1 signal were measured and used for further analysis. On average 40-100 cells were measured for every experimental condition per biological replicate. Three biological replicates were used for each experiment.

#### Sarcomere length

Sarcomere length was measured using combination of Fiji ^40^ and custom R application on 63x objective acquired images. First intensity line profiles along one to four myofibril segments containing ≥ 3 Z-discs per cell were selected. The intensity values and region of interest snapshots were saved in csv and tif format respectively for future validation. Number and spacing of peaks were measured using R based application (available at https://vinarsky.shinyapps.io/sarcomere/). The quality of myofibril segments and Z-disc detection was independently validated by another group member. Fifteen to thirty cells were measured per condition in each biological replicate. Three biological replicates were used for each experiment. In YAP1 re-expression experiments the nuclear intensity of YAP1 was used to identify YAP1 rescued cardiomyocytes.

#### Differentiation efficiency

Differentiation efficiency was measured in single cell cultures of cardiomyocytes. Twenty-seven independent view fields of DAPI and cardiomyocyte markers (cardiac troponin/sarcomeric actinin) were acquired using confocal microscope at 10x objective per experimental condition. After pre-processing in Ilastik pipeline, percecustom Cell Profiler 4 pipeline DAPI channel was used to segment nuclei, cardiomyocyte marker channel was used to identify regions covered by cardiomyocytes. The percentage of cardiomyocytes was calculated as a ratio of nuclei found in cardiomyocytes covered area to total number of nuclei.

#### RNA isolation, reverse transcription PCR and Real-Time PCR

For RNA isolation, iPSC-CMs at day 40 of differentiation were dissociated with MACS Multi Tissue Dissociation Kit 3 (Miltenyi Biotec) according to manufacturer’s instructions. WT and YAP1-KO EHTs were snap frozen in liquid nitrogen and stored at −80 °C before thawing them O/N at −20 °C in RNA LATER-ICE (ThermoFisher). RNA was extracted using High Pure RNA Isolation Kit (Roche) according to manufacturer’s instructions and quantified with Nanodrop 2000 Spectrophotometer (Thermo Fisher Scientific).

RNA reverse transcription was performed with the Transcriptor First Strand cDNA Synthesis Kit (Roche) according to manufacturer’s instructions. One ug of total RNA was used for each sample. RT-PCR was performed using StepOnePlus System (ThermoFisher) with qPCRBIO SyGreen® Mix Hi-ROX (PCR Biosystems) with respective primers (Supplementary Table 2).

### ChIP-seq

Spontaneously contracting confluent cultures of iPSCs derived cardiomyocytes were differentiated in 60 mm tissue culture plates (one plate per sample) until day sixteen of differentiation. Chromatin was immunoprecipitated from three technical replicates using a ChIPgrade anti-YAP antibody (CST14074, Cell Signaling Technologies) and following the manufacturer’s protocol (Pierce™ Agarose ChIP Kit, Thermo Fisher Scientific). A control ChIP was performed using rabbit immunoglobulins (IgG). The samples were eluted in 30 μL eluting buffer and stored at −80 °C before analysis.

For library preparation, the size distribution of each ChIP DNA sample was assessed by running a 1 µL aliquot on Agilent High Sensitivity DNA chip using an Agilent Technologies 2100 Bioanalyzer (Agilent Technologies, Santa Clara, CA, USA). The concentration of each DNA sample was determined using a high sensitivity Quant-iT™ dsDNA Assay Kit and a Qubit Fluorometer (Thermo Fisher Scientific). Purified ChIP DNA (10 µg) was used as the starting material for sequencing libraries preparation. Indexed libraries were prepared using a TruSeq ChIP Sample Prep Kit (Illumina Inc., San Diego, CA, USA). The libraries were sequenced (single read, 1x50 cycles) at a concentration of 10 pm/lane on a HiSeq 2500 (Illumina Inc.).

Data analysis was performed by Genomix4Life S.r.l. (Salerno, Italy). The raw sequence files generated (.fastq) underwent quality control analysis using FastQC (http://www.bioinformatics.babraham.ac.uk). The reads were aligned to the human genome (assembly hg19) using bowtie ^41^, allowing up to one mismatch and considering uniquely mappable reads. The reads of replicates and corresponding input samples were merged for peaks calling as previously described ^42^. ChIP-Seq peaks were identified and analysed using HOMER Motif Database (-F: 2.0, -L: 2.0 and -C: 1.0) with a false discovery rate < 0.01 ^43^. The assignment of YAP peaks to target genes was obtained using the web tool ChIPSeek ^44^.

Through this step, it was possible to assign peaks to the transcription start site (by default defined from - 1 kb to + 100 bp), transcription termination site (by default defined from - 100 bp to + 1 kb), Exon (Coding), 5’-untranslated region (UTR) Exon, 3’ UTR Exon, Intronic or Intergenic regions. As some annotations overlap, the following order of priority was chosen for the assignment:

● Transcription start site (by default defined from - 1 kb to + 100 bp)
● Transcription termination site (by default defined from -100 bp to + 1 kb)
● CDS exons
● 5’ UTR exons
● 3’ UTR exons
● **CpG islands
● **Repeats
● Introns
● Intergenic

Over-represented sequence motifs were defined according to motif descriptors in the JASPAR database and computed using PScan-ChIP ^45^. The following parameters were set:

● Organism: Homo Sapiens Assembly: hg19
● Background: Mixed
● Descriptors: Jaspar 2016

Nucleotide best occurrence was calculated by WebLogo (http://weblogo.berkeley.edu/) by running a Report Best Occurrences analysis on any given transcription factor.

### RNA-seq and data analysis

Spontaneously contracting confluent cultures of iPSCs derived cardiomyocytes were differentiated in 60 mm tissue culture plates (one plate per sample) until day sixteen of differentiation.

Library was prepared using NEBNext® Ultra™ II Directional RNA Library Prep Kit for Illumina® with NEBNext® Poly(A) mRNA Magnetic Isolation Module and NEBNext® Multiplex Oligos for Illumina® (Dual Index Primers Set 1). Kits were employed according to manufacturers’ protocol, input for library preparation was 200-300 ng total RNA.

Sequencing was done on Illumina NextSeq 500 using NextSeq 500/550 High Output v2 kit (75 cycles). We have done single-end 75bp sequencing in multiple sequencing runs until all samples had at least 30 million passing filter reads. Fastq files were generated using bcl2fastq software without any trimming.

The quality of the raw sequencing data was assessed using FastQC (https://www.bioinformatics.babraham.ac.uk/projects/fastqc/) and aligned to the hg38 reference genome using the TopHat2 aligner ^46^. Raw gene counts were obtained by calculating reads mapping to exons and summarized by genes using reference gene annotation (Ensembl 90; Homo sapiens GRCh38.p10, GTF) by HTSeq ^47^.

Differential gene expression was performed using DESeq2 bioconductor package. Genes were considered as differentially expressed when the Benjamini-Hochberg adjusted *P* value ≤ 0.05 and log2 fold-change (log2FC) ≥ 1.5.

The Gene Ontology (GO) categories were downloaded from AmiGO 2 repository (https://amigo.geneontology.org).

### Electrophysiology

After 48 hours, dissociated CMs were recorded at physiological temperature (36 ± 1°C) using the patch-clamp technique in I-clamp or V-clamp mode in whole-cell configuration.

The amplifier Axopatch 200B, the Digidata 1550, and the pClamp 10.0 software (Molecular Devices, LLC) were used for data collecting. 10 kHz sampling and 1–5 kHz filtering were applied to the data. Clampfit 10.0 (Molecular Devices, LLC) in conjunction with Origin Pro 9 (OriginLab) was used to analyse the data.

Spontaneous APs were recorded in CMs superfused with Tyrode solution (pH 7.4) containing (mM): 137 NaCl, 5 KCl, 2 CaCl_2_, 1 MgCl_2_, 10 D-glucose, 10 Hepes–NaOH. Patch pipettes were filled with the intracellular-like solution containing (mM): 120 KCl, 20 Na– HEPES, 10 MgATP, 0.1 EGTA–KOH, and 2 MgCl_2_ (pH 7.1). The following AP parameters were analyzed: rate (Hz), maximum diastolic potential (MDP, mV), AP duration at 50% of repolarization (APD50).

To record the funny current (If), the extracellular solution contained (mM) 110 NaCl, 0.5 MgCl_2_, 1.8 CaCl_2_, 5 Hepes-NaOH, 30 KCl, supplemented with 1 BaCl2 and 2 MnCl2. If was activated from a holding potential (hp) of −30 mV applying 10 mV hyperpolarizing voltage steps to the range of −35/-125 mV long enough to reach steady-state of activation, followed by a fully activating step at -125 mV.

T-type Ca^2+^ current (ICaT) was isolated in Na^+^-free (replaced with equimolar TEA Cl) Tyrode solution, containing 5 mM CaCl_2_ and supplemented with nifedipine (0.01 mM), TTX (0.01 mM) and 4-aminopyridine (2 mM) to block L-type Ca^2+^ channels, Na^+^ channels and K^+^ channels respectively; K^+^ ions in the pipette solution were replaced by equimolar CsCl and TEACl. ICaT was isolated as Ni^2+^ (0.1 mM)-sensitive current, applying depolarizing voltage steps to −20 mV (hp of −80 mV).

L-type Ca^2+^ current (ICaL) was recorded superfusing the cells with (mM) 135 NaCl, 10 CsCl, 1 CaCl_2_, 1 MgCl^2^, 5 HEPES, 10 glucose, supplemented with 0.01 tetrodotoxin (TTX). The intracellular solution contained (mM): 135 CsCl, 1 MgCl^2^, 4 ATP (sodium salt), 0,1 GTP (sodium salt), 5 EGTA–KOH, 5 HEPES-KOH (pH 7.2). ICaL was recorded as nifedipine (0.01 mM)-sensitive current, applying depolarizing voltage steps, 10 mV each, to the range of −40/ +50 mV, followed by a step at 0 mV (hp of −50 mV).

To dissect Na^+^ current (INa), Tyrode solution was supplemented with 0,01 mM nifedipine. INa was recorded as TTX (0.03 mM)-sensitive current, applying depolarizing voltage steps, 10 mV each, to the range of −80/ +60 mV, followed by a step at −20 mV (hp of -90 mV).

The I_NCX_ was evaluated as the Ni^2+^ (10 mM)-sensitive current during voltage ramps (100 mV/s) from +60 mV to −100 mV (hp −40 mV). The external solution contained (mM): 135 NaCl, 10 CsCl, 1 CaCl_2_, 1 MgCl_2_, 10 Hepes-NaOH, 10 TEA-Cl, 10 D-glucose (pH 7.35) to which 0.2 mM BaCl2, 0.005 mM nifedipine, 0.05 mM lidocaine, and 1 mM ouabain were added to block K^+^, Ca^2+^, Na^+^ channels, and the Na^+^/K^+^ pump, respectively. Pipette was filled with (mM): 140 CsOH, 75 aspartic acid, 20 NaCl, 10 CaCl^2^, 10 HEPES, 20 EGTA, 20 TEA-Cl, 5 MgATP (pH 7.3). I_NCX_ density at −80 mV and + 40 mV was taken as representative of the forward and the reverse mode working direction of NCX, respectively.

Current densities were calculated by measuring in each cell the membrane capacitance (C_m_). The steady-state activation and inactivation curves were obtained from normalized tail currents or conductances and interpolated with the Boltzmann equation.

The analyses were carried out in parallel between WT and YAP1-KO derived hESC-CMs (N ≥ 3). Significant alterations have been delineated between the two groups by *P* < 0.05 with Student’s *t* test comparison after checking for normal distribution of data with the Shapiro–Wilk test.

### Intracellular Ca^2+^ dynamics

Cytosolic Ca2+ was recorded in single V-clamped CMs loaded with Fluo4-AM (10 µM) at physiological temperature as previously described ^48^. Cells were incubated for 30 min with Fluo4 at room temperature and then the dye was washed for at least 10 min to allow its de-esterification. Fluo4 was excited at 488 nm, and the emission was collected through a 530 nm band-pass filter, converted to voltage, low-pass filtered (200 Hz) and digitized at 2 kHz after further low-pass digital filtering (FFT, 50 Hz).

Cells were superfused at physiological temperature with the Tyrode’s solution (mM) containing: 154 NaCl, 4 KCl, 2 CaCl_2_, 1 MgCl_2_, 5 HEPES/NaOH, and 5.5 D-glucose (pH 7.35) to which 1 mM BaCl2 and 2 mM 4-aminopyridine were added to block K^+^ channels. Patch-clamp pipettes were filled with (mM) 23 KCl, 110 KAsp, 0.04 CaCl_2_, 3 MgCl_2_, 5 HEPES-KOH, 0.1 EGTA-KOH, 0.4 NaGTP, 5 Na2ATP, 5 Na2-phosphocreatine and 0.01 mM Fluo4-K^+^ salt (pH 7.3). Ca^2+^ transients (CaT) were evocated at 0.5 Hz during 100 ms steps to 0 mV after 50 ms step to −35 mV to inactivate Na^+^ channels. Sarcoplasmic reticulum (SR) Ca^2+^ content (CaSR) was estimated at steady state by measuring CaT amplitude elicited by electronically timed (10 mM) caffeine pulse after 10s at −80 mV. Fluo4 signal was normalized to the resting fluorescence (F0) before caffeine superfusion.

## Statistical analyses

All data are expressed as means± SD unless otherwise specified. The statistical analyses were performed by using GraphPadPrism 8.0 software or R. The null hypothesis was rejected when *P*<0.05. The number of biological (N) and technical (n) replicates and the statistical tests used are indicated in figure legends and in data files. Electrophysiological data and Ca^2+^ dynamics are shown as mean ± SEM

## Results

### YAP1 acts as a transcriptional activator in hESC-CMs

To understand YAP1’s role in the cellular context of CM maturation and function we first investigated the effects of YAP1 deficiency (KO) on the transcriptional landscape of hESC-CMs. For this purpose, YAP1-KO hESCs and their isogenic control line (hereafter referred to as WT) were differentiated into spontaneously beating CMs according to an established protocol^32^ for 15 days. At this time-point, total RNA was harvested and analyzed by RNA sequencing. Principal component analysis (PCA) of the RNA-seq data indicated that YAP1 depletion consistently and reproducibly shifted hESC-CMs transcriptional landscape away from the WT. One quarter (4473 out of 17499) of all detected genes were differentially regulated in YAP1-KO CMs (fold change >1.5 and adjusted p. value <0.05) (Figure 1A; Supplementary Table 3). Importantly, genes involved in late CM maturation (MYOZ2, EMILIN2, MYH7, TNNT1, ACTN3) were significantly downregulated in YAP1-KO cells, which expressed significantly higher levels of early cardiac commitment markers (ACTA1, NKX2-5, MEF2C, ISL1, ACTA2) (Figure 1B).

**Figure 1:**
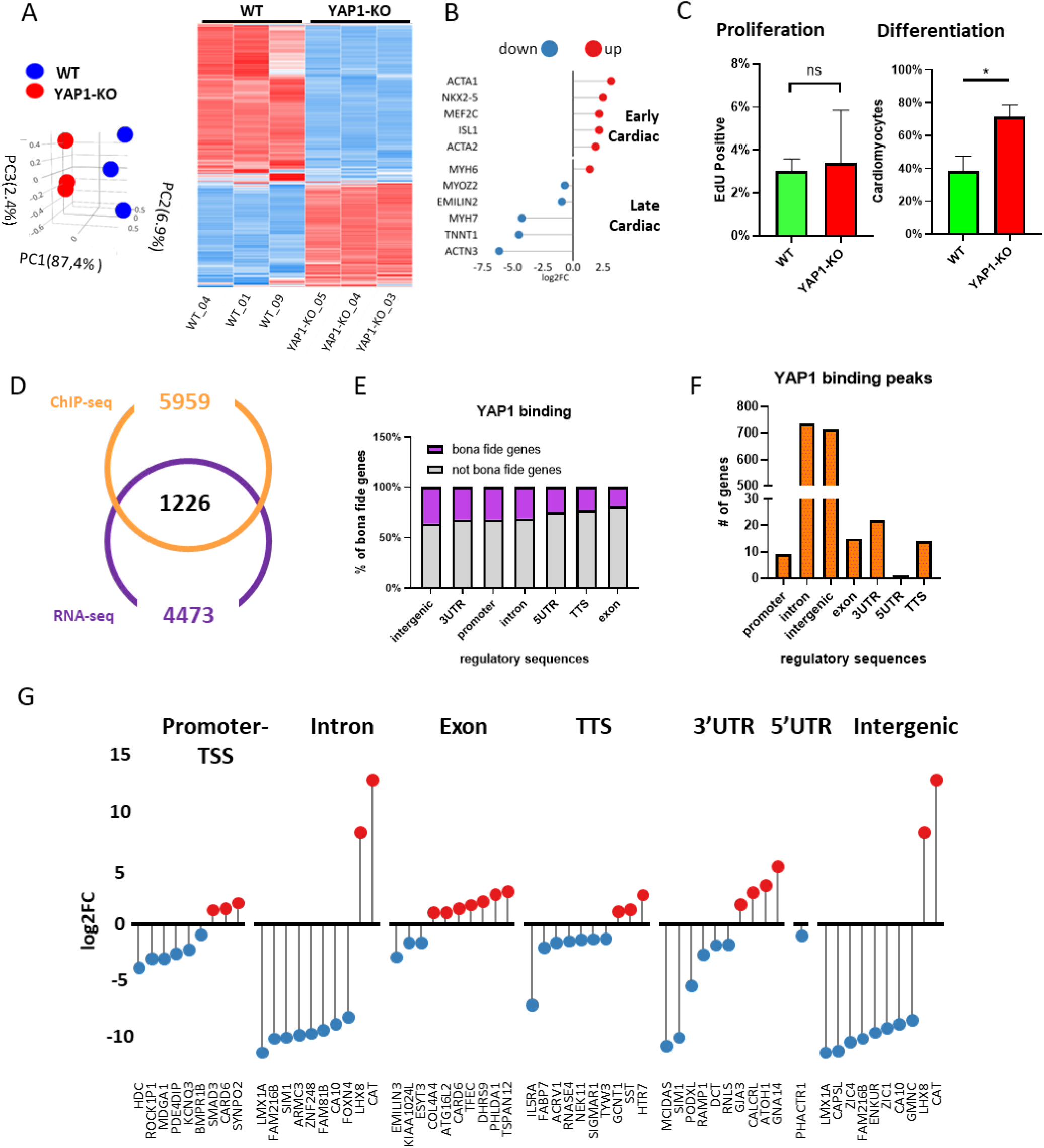
YAP1 transcriptional activity in hESC-CMs. Principal component analysis (PCA) (left) and heatmap representation (right) of the differentially expressed genes (DEG) in WT and YAP1-KO hESC-CMs as identified by RNA-sequencing (day 15 of differentiation, n=3) (A). Lollipop plot representation of differential gene expression of key CM maturation genes in YAP1-KO compared to WT hESC-CMs (n=3) (B). Bar plot representation of percentage of EdU-positive (left) and troponin-T positive (right) YAP1-KO and WT hESC- CMs (n=3) (C). Venn diagram representation of YAP1 *bona fide* targets, as the genes both identified as YAP1 targets in ChIP-seq analysis and differentially expressed in YAP1-KO RNA-seq (D). Bar plot representation of percentage of *bona fide* genes with YAP1 peaks in different genomic regulatory sequences (E). Bar plot representation of frequency of YAP1 binding to indicated regulatory genomic sequences (F). Lollipop plot representation of gene expression of the top 10 differentially regulated *bona fide* genes grouped by genomic regulatory sequences (G).

Altogether these results indicated that, although hESCs in which YAP1 has been genetically depleted retain the ability to differentiate into early contractile cells, the absence of the protein affects their maturation. This result could not be explained by a difference in the proliferation of less mature YAP1-KO hESC-CMs (3.0%±0.6% in WT and 3.4%±2.5% in YAP1-KO) (Figure1C, left), but rather by their differentiation efficiency (38,5%± 9.06% in WT vs 71.44%± 7,30% in YAP1-KO) (Figure 1C, right).

To investigate how YAP1 depletion affected hESC-CMs transcriptional profile, we immunoprecipitated endogenous YAP1 protein in WT differentiated cells and analyzed YAP1 DNA binding activity by chromatin immunoprecipitation (ChIP) followed by DNA sequencing (seq). See Supplementary Table 4 for the complete list of YAP1 binding sites in hESC-CMs. We integrated the data obtained by RNA-seq in YAP1-KO hESC-CMs with those generated by ChIP-seq analysis in the WT counterpart to identify YAP1 *bona fide* targets as those genes which are physically bound by the protein and changed their expression. We found that 27.4% (1226 out of 4473) of the genes differentially regulated were *bona fide* targets (Figure 1D). The highest frequency of *bona fide* target genes was associated with YAP1 binding to intergenic sequences (36%), lowest with YAP1 binding to exons (19%) (Figure 1E). The highest proportion of YAP1 binding peaks in sequences of *bona fide* targets was found in introns and intergenic regions which accounted for over 700 genes each (734 intron bound, 714 intergenic bound) (Figure 1F). The prevailing effect of YAP1 binding on genomic regulatory sites was determined by quantification of *bona fide* gene expression in YAP1 deficient cardiomyocytes. Six out of nine *bona fide* genes where YAP1 was bound to promoter sequences (TSS) were repressed in YAP1-KO CMs. Similarly, out of the top ten differentially expressed genes, where YAP1 binding was detected in introns, eight genes were repressed. The same pattern was observed in *bona fide* genes where YAP1 was bound in transcription termination sequence (TTS), 3’ untranslated region (3’UTR), and intergenic region (Figure 1G). To enlarge the number of *bona fide* targets, we investigated the distribution of YAP1 peaks on enhancers and promoters annotated in GeneHancer ^49^, identifying its presence on 328 enhancers and 120 promoters (*P* value<0.05 with 1000 permutation), that putatively regulates 134 differentially expressed genes. We next browsed through the *bona fide* targets to identify transcription factor regulatory sites. We found a plethora of candidate transcription factors possibly interacting with YAP1 in hESC-CMs which determined the differential expression of the identified genes. Among them, we also detected NKX2-5 and MEF2C which are well known master regulators of CM phenotype (Supplementary Figure 1A). Taken together our data suggest that YAP1 predominantly exerts a positive effect on the transcriptional activity in hESC-CMs.

### YAP1 indirectly activates the transcription of genes involved in sarcomere maturation in hESC-CMs.

To disclose the mechanism through which YAP1 transcriptional activity contributes to hESC-CMs maturation, we focused on gene subsets essential for CM phenotype acquisition and function, and relevant for cardiac muscle development and sarcomere structure. In the absence of a specific annotation among the YAP1 *bona fide* targets that could account for sarcomere-related categories, we looked for indirect targets in the RNA-seq dataset. Among the genes found consistently dysregulated in YAP1-depleted cells, we identified numerous genes involved in the development of muscle structure, sarcomere, and Z-disc. In particular, we confirmed the significant down-regulation of the cardiac mesoderm specification WNT3A ^36, 50^ and observed a similar tendency in sarcomere and Z-disc associated genes (ACTN3, TNNT1, MYH7, CRYAB, IGFN1), genes connected with sarcomere turnover (FBXL22), actin binding (PDLIM2, LMOD2), and Rho-activated transcription (ABRA) (Figure 2A; Supplementary Figure 1B).

**Figure 2:**
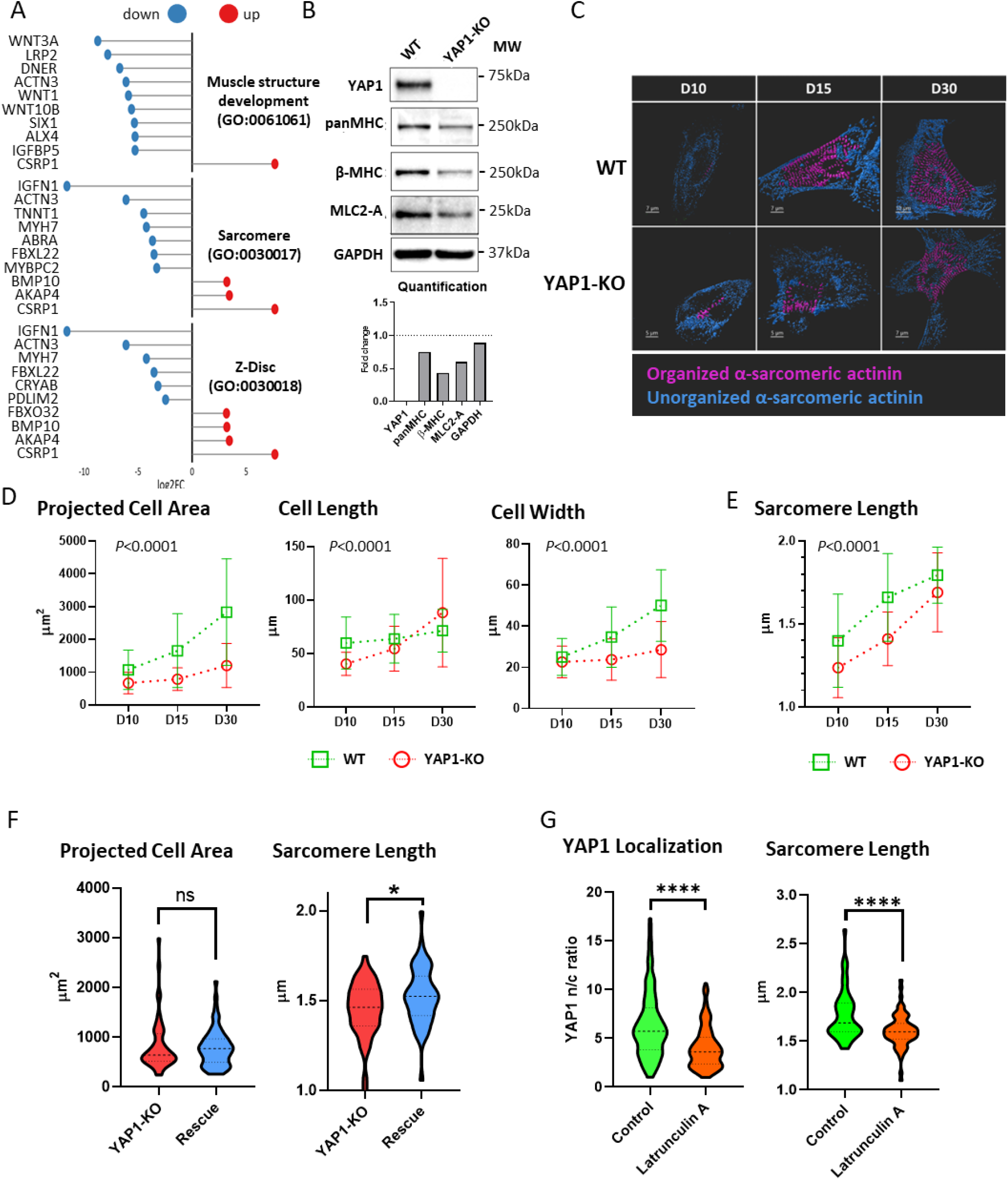
YAP1 is instrumental for sarcomere assembly and maturation in hESC-CMs. Lollipop plot representation of up to 10 most differentially expressed (|Log2FC|) genes of indicated gene ontology (GO) categories in YAP1-KO compared to WT hESC-CMs (n=3) (A). Western blot analysis of the indicated sarcomeric proteins in YAP1-KO and WT CMs quantification on bottom; GAPDH was used for normalization (B). Characterization of changes in morphology and sarcomere length in WT and YAP1-KO CMs at days 10, 15, and 30 of differentiation. Representative segmentation of α-sarcomeric actinin into organized (magenta) and disorganized (blue) structures (C). Morphometry of projected cell area, length, width (D), and sarcomere length (E). Statistics: Two way ANOVA, Interaction: *P*<0.0001. Violin plot representation of projected cell area (left) and sarcomere length (right) of YAP1-KO CMs and YAP1-KO CMs rescued by ectopically expressed YAP1 T2A-mCherry plasmid (Qin et al. 2016) (F). hESC-WT CMs on day 16 were untreated (Control) or incubated with latrunculin A. Analysis of YAP1 localization by nuclear/cytoplasmic (n/c) ratio of YAP1 intensity and sarcomere length (G). The data are shown as mean±S.D. (N=3). Statistics: Unpaired *t* test, ns: *P*>0.05, \**P*<0.05, \*\*\*\**P*<0.0001.

The increased cell size, myosin heavy chain 7 (MYH7) expression, sarcomeric actinin organization into Z-discs, and enhanced sarcomere length are signs of CM maturation both *in vivo* and *in vitro* ^51, 52^. While YAP1 deficiency did not inhibit spontaneous contractions, we noted that bulk cultures of YAP1-KO CMs consistently expressed less sarcomere proteins (Figure 2B). In addition, evident qualitative differences in cell size and sarcomeric actinin organization in YAP1-KO hESC-CMs throughout differentiation (Figure 2C; Supplementary Figure 2A for better resolution) prompted us to investigate CM morphology and contractile apparatus in detail. We turned to super-resolution microscopy to quantify differences in projected cell area, morphology, and sarcomere length from the onset of beating (day 10) to day 30 of differentiation. The projected area of WT CMs increased nearly threefold from 1068± 601 µm^2^ at day 10 to 2831±1625 µm^2^ at day 30 (*P*<0.001, two way ANOVA), with major contribution coming from the cardiomyocyte width (from 25±8.9 µm to 50±17.4 µm). YAP1-KO CMs growth was instead limited to less than twofold (*P*<0.0001, two way ANOVA) with no significant change in CM width (from 23±8 µm at day 10 to 29±13.6 µm at day 30) (Figure 2D). These data indicate that cell growth - and in particular lateral growth - depends on YAP1 activity during *in vitro* differentiation.

Next, we measured sarcomere length in WT and YAP1-KO cardiomyocytes during differentiation. From the beginning of myofibril assembly (at day 10) sarcomere length was markedly shorter in YAP1-KO cells (1.39±0.281 µm in WT vs 1.24±0.18 µm in YAP1-KO; multiple comparison t-test Q=1%, *P*=0.0163). By day 15 the difference in sarcomere lengths further increased; YAP1-KO CMs sarcomeres were on average 0.25µm shorter (1.66±0.264 µm in WT vs 1.41± 0.161 µm in YAP1-KO; multiple comparison *t* test Q=1%, *P*<0.0001). At day 30 the difference in sarcomere length narrowed to 0.1 µm but was still significant (1.79±0.169 µm in WT vs 1.69±0.238 µm in YAP1-KO; multiple comparison *t* test Q=1%, P=0.0004) (Figure 2E).

Observing that the most dramatic differences in the growth of cell area and sarcomere occurred between days 10 and 15 of differentiation prompted us to investigate the effect of YAP1 activity on cardiomyocyte maturation after the onset of beating. First, we transiently expressed full length *YAP1* in YAP1-KO CMs at D12 (rescue). YAP1 rescue had no significant effects on projected cell area (836±532 μm^2^ in WT vs 786.3±379 μm^2^ in rescue; unpaired *t* test, *P*=0.5618) (Figure 2F, left), length, and width (data not shown) after eight days of YAP1 re-expression. Interestingly, we observed partial yet significant recovery of sarcomere length (1.45±0.153 μm in YAP1-KO vs 1.53±0.163 μm in YAP1 rescue; unpaired *t* test, *P*=0.03) (Figure 2F, right). Finally, to further corroborate the effect of YAP1 transcriptional activity on sarcomere length we inhibited YAP1 activity by blocking its nuclear localization by cytoskeletal tension inhibitor latrunculin A, known to impair YAP1 nuclear localization ^9^. Here we observed reduced nuclear localization of YAP1 (YAP1 nucleus/cytoplasm ration: 6.9±4.50 in control CMs vs 4.8±4.01 in latrunculin A treated) (Figure 2G, left) and significant disruption of myofibrillar structures (Supplementary Figure 2B). Interestingly, sarcomere length was significantly reduced by latrunculin A (1.77±0.255 μm in control CMs vs 1.61±0.171 μm in latrunculin A treated; unpaired *t* test, *P*<0.0001) (Figure 2G, right).

Taken together these data indicate that YAP1 is needed for sarcomere assembly and maturation in CMs derived from hESCs.

### YAP1 modulates CMs electrophysiological properties

The process of CM specification and maturation forces dramatic changes in the ability of CMs to initiate and propagate electrical impulses and generate force ^53^. We observed that YAP1 deficiency dysregulated multiple categories of genes modulating CM beating rate and intracellular Ca^2+^ dynamics (Figure 3A; Supplementary Figure 3). Specifically, gene expression of Hyperpolarization-activated cyclic nucleotide–gated (HCNs) channel, and key components of Ca^2+^ homeostasis were downregulated (i.e. CALM2, CAMK2D) in YAP1-KO CMs.

**Figure 3:**
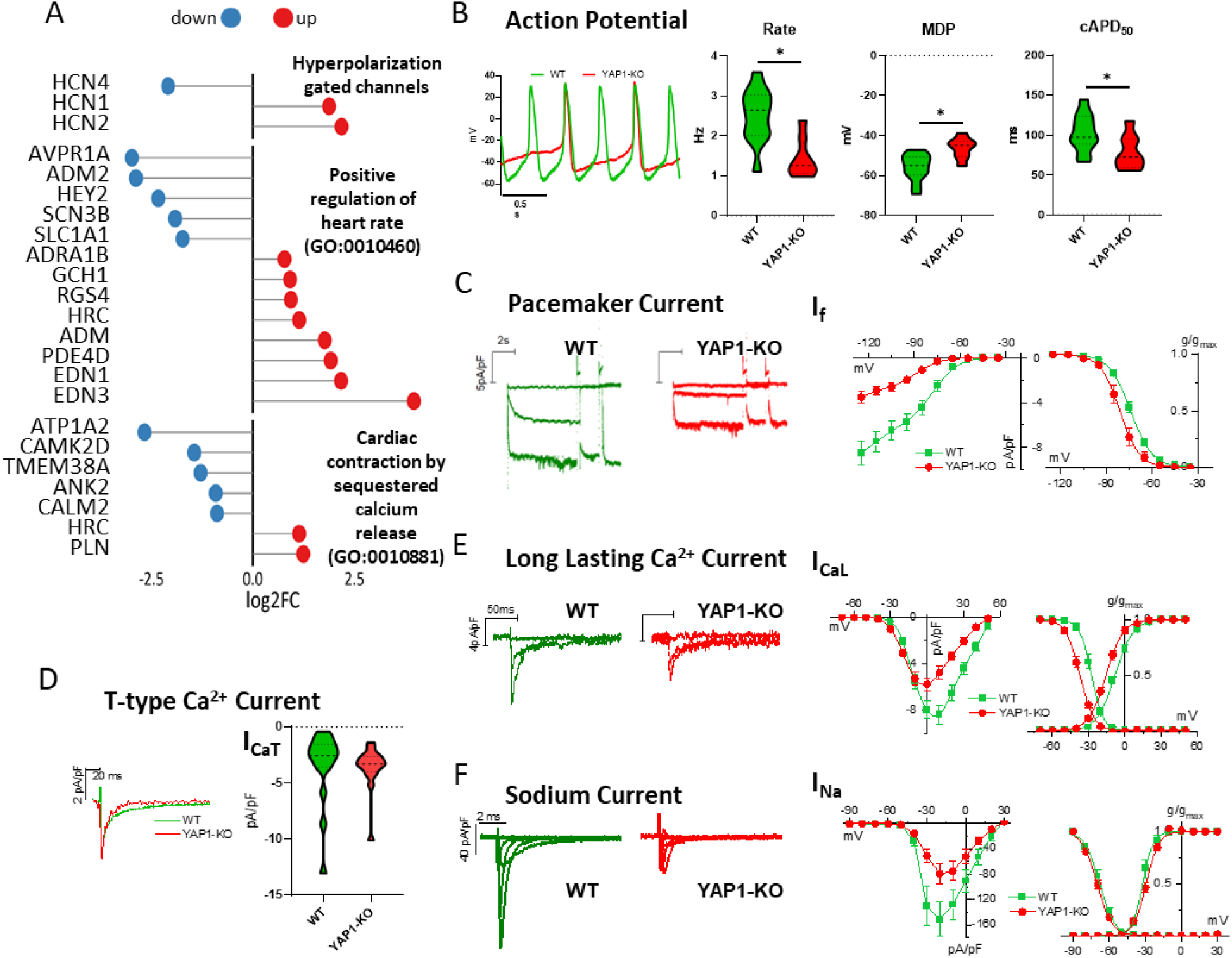
YAP1 regulates maturation of electrophysiological properties of hESC-CMs. Lollipop plot representation of differential gene expression of indicated gene categories in YAP1-KO compared to WT hESC-CMs (n=3) (A). Representative action potentials (B, left) and quantification of AP parameters in WT (n=12) and YAP1-KO (n=8) CMs: beating rate, maximum diastolic potential (MDP), corrected AP duration at 50% of repolarization (cAPD_50_) (B, right). Representative recordings of pacemaker current (I_f_) (C, left), I_f_ I/V relations and steady-state activation curves (left) in WT (n=23) and YAP1-KO (n=14) CMs (C, right). Representative recordings (D, left) and quantification of peak T-type Ca^2+^ current (I_CaT_) at −20 mV in WT (n=16) and YAP1-KO (n=16) CMs (D, right). Representative recordings of long lasting Ca^2+^ current (I_CaL_) (E, left), ICaL I/V relations and steady-state activation/inactivation curves in WT (n=23) and YAP1-KO (n=18) CMs (E, right). Representative recordings of sodium current (INa) (F, left), I_Na_ I/V relations and steady-state activation/inactivation curves in WT (n=12) and YAP1-KO (n=18) CMs (F, right). Statistics: unpaired *t* test, \**P*<0.05.

To extend our knowledge on the importance of YAP1 for CM maturation from a functional perspective, we analyzed the electrophysiological properties and the intracellular Ca^2+^ dynamics of YAP1-KO and WT CMs using patch-clamp and fluorimetric techniques, respectively. At 20 days of differentiation, both YAP1-KO and WT presented spontaneous electrical activity as shown by representative action potential (AP) traces (Figure 3B). In comparison to WT cells, YAP1-KO CMs beat significantly slower, showed more depolarized maximal diastolic potentials (MDP) and shorter AP duration at 50% of repolarization (APD50) (Figure 3B, right). To notice, APD50 was corrected to the cell beating rate (cAPD50) to avoid APD changes mainly related to different beating rates.

To shed light on the AP rate dysregulation, we measured the pacemaker current (I_f_) (Figure 3C, left) dependent on HCNs expression and well known to contribute to diastolic depolarization rate (DDR). We observed that If density was significantly reduced in YAP1- KO CMs (Figure 3C, right). Moreover, the steady-state activation curve was significantly leftward shifted (toward more negative potentials, −7,08±0,17 mV, *P*<0.05) (Figure 3C, right), suggesting that both maximal conductance reduction and changes in activation biophysics of If could explain the decreased beating rate of YAP1-KO CMs. Notably, T-type Ca^2+^ current (ICaT), potentially contributing to DDR, was not affected by YAP1 deficiency (Figure 3D).

According to APD50 shortening, the Long lasting Ca^2+^ current (I_CaL_) was significantly reduced in YAP1-KO CMs in terms of conductance and biophysics (Figure 3E, left). Indeed, voltage-dependent ICaL availability (Figure 3E, right) was also affected by YAP1 deficiency; in particular, both steady-state activation and inactivation curves were leftward shifted in YAP1-KO CMs of −7,4±0,2 mV (*P*<0.05 vs WT) and −9,62±0,17 mV (*P*<0.05 vs WT) respectively (Figure 3E, right), suggesting a marked alteration of ICaL voltage dependency in YAP1-KO CMs.

Finally, Na^+^ current density (INa) was markedly reduced in YAP1-KO CMs in comparison to WT (Figure 3F, left), while its activation/inactivation properties were unaffected by the lack of YAP1 (Figure 3F, right).

### YAP1 modulates CMs intracellular Ca^2+^ dynamics

Analogously to electrophysiological properties, Ca^2+^ dynamics also change during CM development and maturation ^53^. To study how YAP1 genetical depletion affects Ca^2+^ handling properties in cardiomyocytes, we evaluated YAP1 effects on voltage- and caffeine-induced Ca^2+^ transients (CaT) in voltage-clamped YAP1-KO and WT CMs.

We controlled the membrane potential to avoid confounding secondary effects otherwise present in spontaneous beating or field stimulated cells. As shown in Figure 4A, compared to WT CMs, YAP1-KO CMs showed slower voltage-induced CaT onset (quantified as increased CaT time to peak, TTP) (Figure 4A, left and middle), and slower CaT decay, suggesting alterations in EC-coupling machinery and in diastolic Ca^2+^ removal systems. Moreover, CaT amplitude tended to be reduced in YAP1-KO CMs (p>0.05 vs WT). In addition, quantification of CaT decay components revealed that the slow component of the CaT decay (quantified as t50 and t90) significantly increased in YAP1-KO CMs, while the early phase of CaT decay (quantified as t20) was unaffected (Figure 4A, right). To better estimate the potential effects of YAP1 on sarcoplasmic reticulum (SR) Ca^2+^ content (CaSR), caffeine-induced CaT and the Na^+^/Ca^2+^ exchanger (NCX) current (I_NCX_) activated during caffeine pulse, were quantified at the same time. Both caffeine-induced CaT amplitude (Figure 4A, left and middle) and Ca^2+^ influx through NCX (not shown, see Methods) were significantly reduced in YAP1-KO CMs, thus indicating a reduced CaSR in comparison to WT cells.

**Figure 4:**
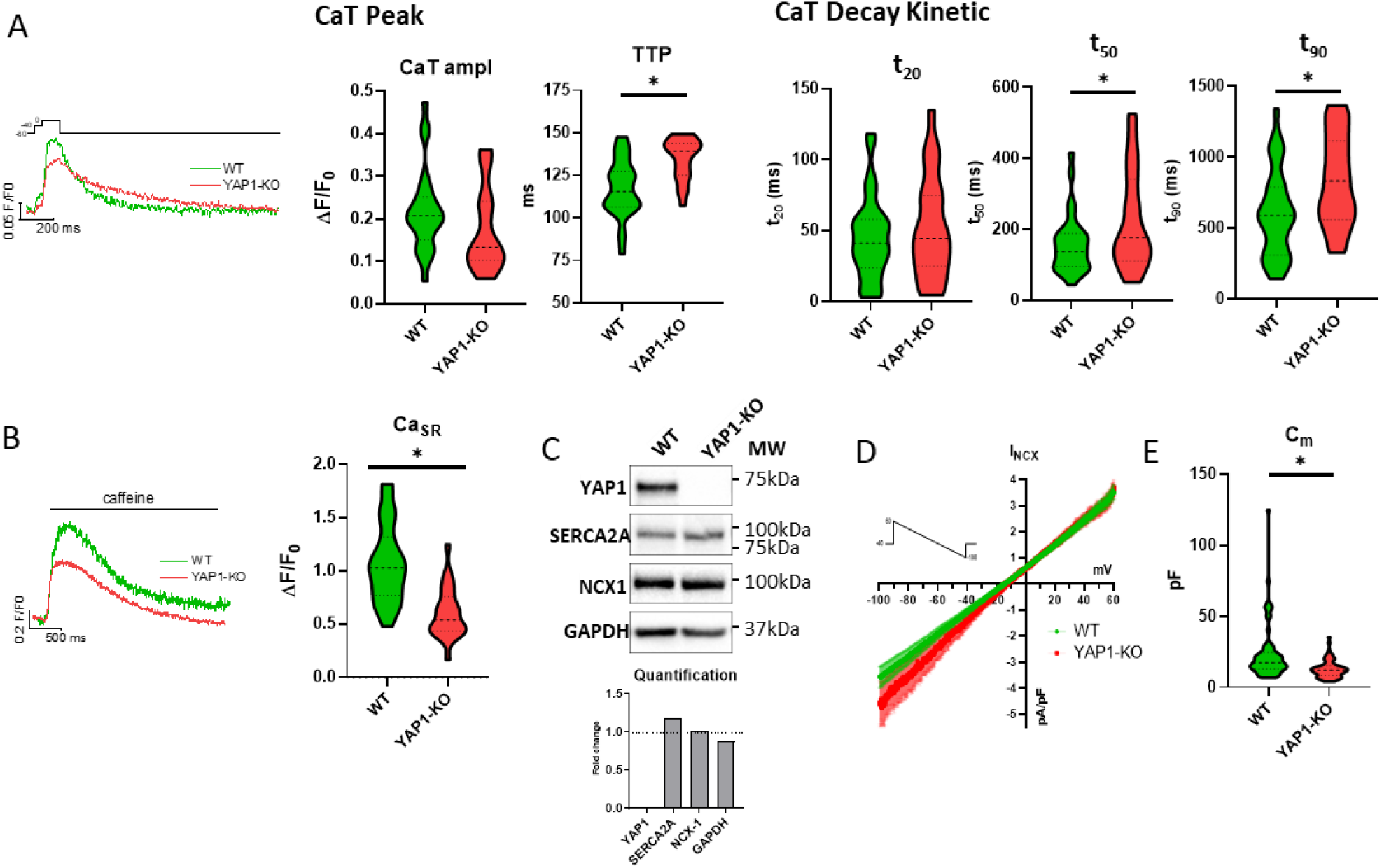
YAP1 regulates Ca^2+^ dynamic of hESC-CMs. Representative traces of voltage-induced Ca^2+^ transient (CaT) (A, left) and quantification of CaT parameters: time to peak (TTP), amplitude, and decay kinetics at 20%, 50%, 90% decay time in WT (n=29) and YAP1-KO (n=23) CMs (A, right). Representative traces of caffeine-induced CaT (B, left) and quantification of the peak amplitude (i.e the SR Ca^2+^ content, CaSR), in WT (n=24) and YAP1-KO (n=22) CMs (B, right). Western blot analysis of SERCA2a and NCX1 protein levels (quantification on bottom) (C). Mean (± sem) I/V relations of NCX current (I_NCX_) in WT (n=25) and YAP1-KO (n=24) CMs (D). Statistics of cell membrane capacitance (C_m_) in WT (n=62) and YAP1-KO (n=66) CMs (E). Statistics: unpaired *t* test, \**P*<0.05.

Furthermore, western blot evaluation of total NCX1 and SERCA2a protein levels did not highlight changes in YAP1-KO CMs in comparison to WT (Figure 4C). To further characterize this finding, we also evaluated NCX1 protein expression at the membrane level by measuring I_NCX_ through a dedicated voltage clamp protocol. Consistent with our western blot results, we did not detect any difference in inward and outward conductance of NCX1 in YAP1-KO CMs compared to WT (Figure 4D). These results are apparently in contrast to the observed slower CaT decay in YAP1-KO CMs and implied an altered distribution of NCX probably related to more fetal-like phenotype of YAP1-KO CMs, leading to a less efficient EC-coupling. Electrophysiological quantification of cell dimension through cell membrane capacitance (C_m_) evaluation confirmed the immature/smaller phenotype of YAP1-KO CMs compared to WT (Figure 4E).

In conclusion, all these measurements point to altered maturation of excitation-contraction coupling mechanisms in absence of YAP1.

### YAP1 is required for compensatory cardiomyocyte growth response to stimuli

Having established the importance of YAP1 transcriptional control over the maturation of sarcomere structure and CM functional properties, we decided to investigate the role of YAP1 by establishing a reductionist *in vitro* model of the myocardial infarction (MI) border zone. The conditions in MI border zone are distinct from the rest of the heart in composition and mechanical forces. CMs at the border of the MI are surrounded by ECM characterized by increased amount of fibronectin and altered strain ^54, 55^. As previously mentioned, nuclear YAP1 localization and distinct transcription profile were reported in CMs in MI border zone *in vivo* ^20, 56^. To create MI border zone-like model of fibronectin deposition and increased strain *in vitro* we exposed hESC-derived cardiomyocytes cultured onto 10 kPa polydimethylsiloxane (PDMS) coated with 1 μg/mL or 10 μg/mL of fibronectin to mechanical actuation (120% static stretch for 24 hours).

As expected, fibronectin deposition induced the nuclear localization of YAP1 in WT CMs (nuclear/cytoplasm ratio 2.52±0.731 in 1 μg/mL vs 4.83±2.357 in 10 μg/mL, unpaired *t* test, *P*<0.0001) (Figure 5A, left). Static stretch also induced the translocation of YAP1 into CM nucleus (from 4.46±2.840 in control vs 7.835±5.565 actuated, unpaired *t* test, *P*<0.0001) (Figure 5A, right). Having established the ability of MI border zone-like conditions to induce the nuclear localization of YAP1 in WT hESC-CMs we focused on the role of YAP1 activity activity in controlling stress-dependent CM hypertrophic growth. Fibronectin deposition increased the projected area of both WT and YAP1-KO CMs (from 564±302.6 µm^2^ at 1 μg/mL to 1454±840.2 µm^2^ at 10 μg/mL, *P*<0.0001 in WT CMs, and from 297±100.6 µm^2^ at 1μg/mL to 518±239.4 µm^2^ at 10 μg/mL *P*<0.0001 in YAP1-KO) (Figure 5B). Remarkably, in all conditions YAP1-KO cells were significantly smaller than the WT counterparts.

**Figure 5:**
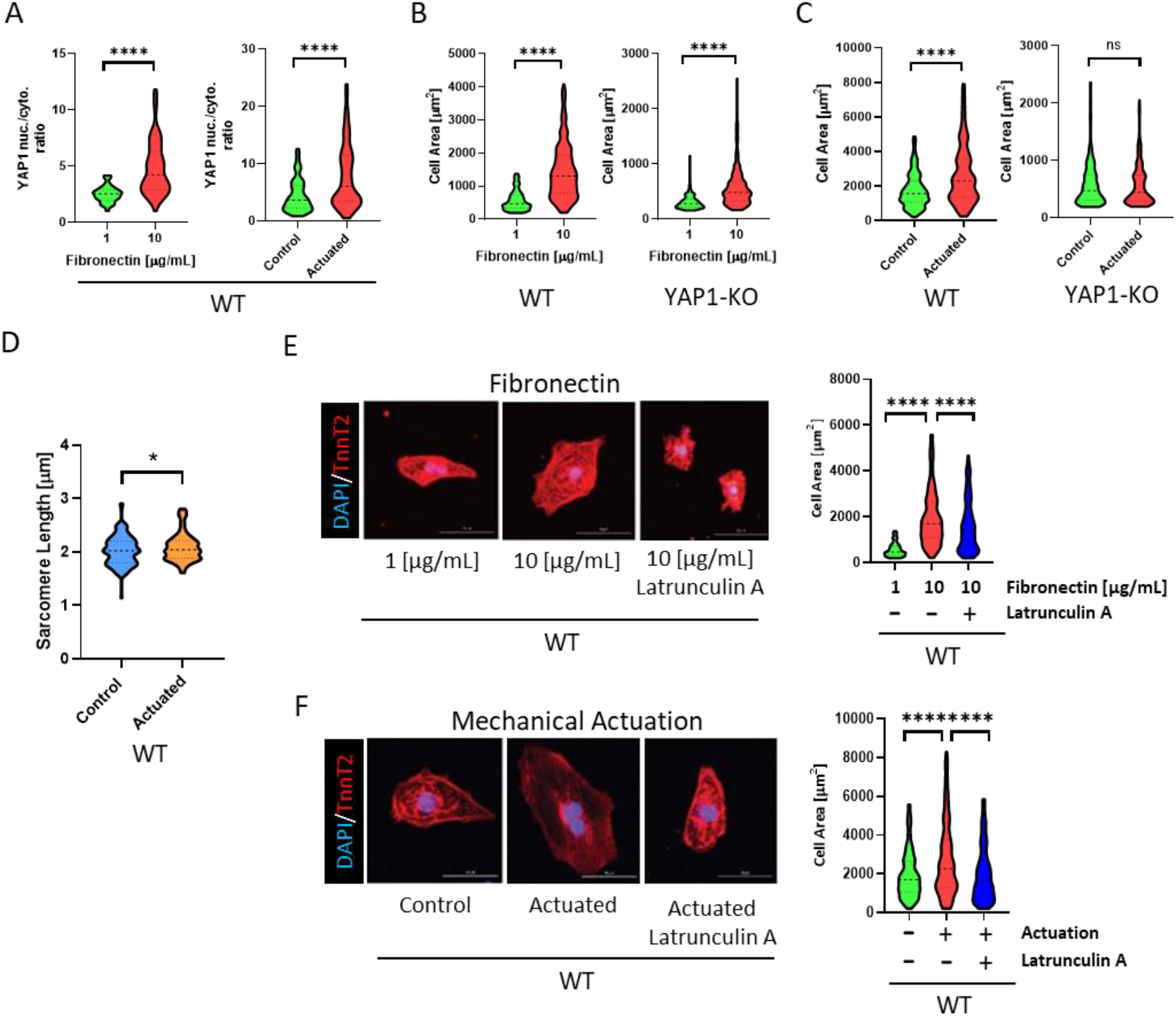
YAP1 activity is required for compensatory growth. Violin plot representation of nuclear translocation of YAP1 in hESC-WT cardiomyocytes in response to fibronectin (left) (*P*<0.0001, N=3, n=145) and mechanical actuation (right) (*P*<0.0001, N=3, n=502) (A). Violin plot representation of projected cell area in WT and YAP1-KO CMs in response to fibronectin (*P*<0.001, N=3, n=253) (B) and mechanical actuation (*P*<0.001, N=3, n=264) (C). Violin plot representation of sarcomere length in response to mechanical actuation in hESC-WT CMs (*P*=0.04, N=3, n=122) (D). Attenuation of cellular response in presence of latrunculin A mediated cytoskeletal tension inhibition in response to fibronectin (E), mechanical actuation (F) in hESC-WT CMs. Representative confocal images of hESC-WT CMs stained for cardiac troponin-T (red) and DAPI (blue), images (E and F, left), quantification of projected cell area upon fibronectin deposition (*P*<0.0001, N=3, n=215) (E, right) and mechanical actuation (*P*<0.0001, N=3, n=328) (F, right). Statistics: Unpaired *t* test, ns: *P*>0.05, \**P*<0.05, \*\**P*<0.01, \*\*\**P*<0.001, \*\*\*\**P*<0.0001.

Static stretch (120%, 24 hours) increased cell area only in WT CMs (1760±991.7 µm^2^ control, 2667±1710 µm^2^ actuated, unpaired *t* test, *P*<0.0001); this response was totally abolished in YAP1-KO CMs (568±349.5 µm^2^ control vs 570±8 µm^2^ actuated, unpaired *t* test, *P*=0.9267) (Figure 5C). It is of note that while the projected area of WT cardiomyocytes increased nearly twofold, sarcomere length increase was modest – while statistically detectable 0.065±0.031 µm (*P*=0.036, unpaired *t* test). Therefore, sarcomere length is not affected by static stretch (Figure 5D).

To confirm the direct role of YAP1 transcriptional activity in the model of fibronectin deposition and increased strain, we reduced cytoskeletal tension and YAP1 nuclear localization in WT CMs by latrunculin A. While we observed statistically significant abolition of compensatory cell growth in both fibronectin deposition and increased strain conditions, the effects of latrunculin A were more pronounced in the strain induced compensatory growth (Figure 5E and 5F). This data is congruent with our observations of partial dependence of compensatory growth on YAP1 activity in response to fibronectin deposition and total YAP1 dependence on mechanically induced compensatory growth.

Taken together these data indicated that YAP1 is required for the compensatory response to our model of pathological stimuli. This response is more pronounced upon mechanical stimulation.

### YAP1 promotes contractile force in 3D microtissue model

Our observation that YAP1 activity is required for sarcomere assembly and intracellular Ca^2+^ dynamics pointed to net positive effect of YAP1 on cardiomyocyte contractility. To test our hypothesis we used a different YAP1^-/-^ cell line ^31^ and more physiologically relevant 3D model of engineered heart tissues (EHT) leveraging recently developed platform to measure contractile force in real time (Cuore, Optics11life) ^35^. Control and YAP1-KO iPSCs differentiated into cardiomyocytes for thirty days, were either kept in 2D or cast into a collagen hydrogel and followed for ten days in a cultivation platform enabling live monitoring of force generation.

First, to verify effects on cardiomyocyte maturation in absence of YAP1 in a different cell line we differentiated cardiomyocytes from control (WT) and YAP1 deficient (YAP1-KO) human induced pluripotent stem cells (iPSCs) ^31^. To assess maturation state we compared ratio of expression of genes known to change with maturation ^57^: myosin heavy chains (MYH7/MYH6), myosin light chains (MYL2/MYL7), and troponin-I isoforms (TNNI3/TNNI1). Indeed, we observed marked reduction in all ratios in YAP1-KO hiPSC-CMs in both 2D and 3D culture confirming impaired maturation in absence of YAP1 (Figure 6A). To further characterize the 3D model, we analyzed myofibril structure in EHTs generated from control and YAP1 deficient cardiomyocytes. As expected, the myofibril content was visibly reduced in EHTs obtained from YAP1 deficient cardiomyocytes (Figure 6B). In addition, the maturation of sarcomere in the YAP1 deficient 3D constructs was significantly compromised as measured by sarcomere length (1.918±0.459 μm in WT vs 1.166±0.353 μm in YAP1-KO, unpaired *t* test, *P*<0.0001) (Figure 6C).

**Figure 6:**
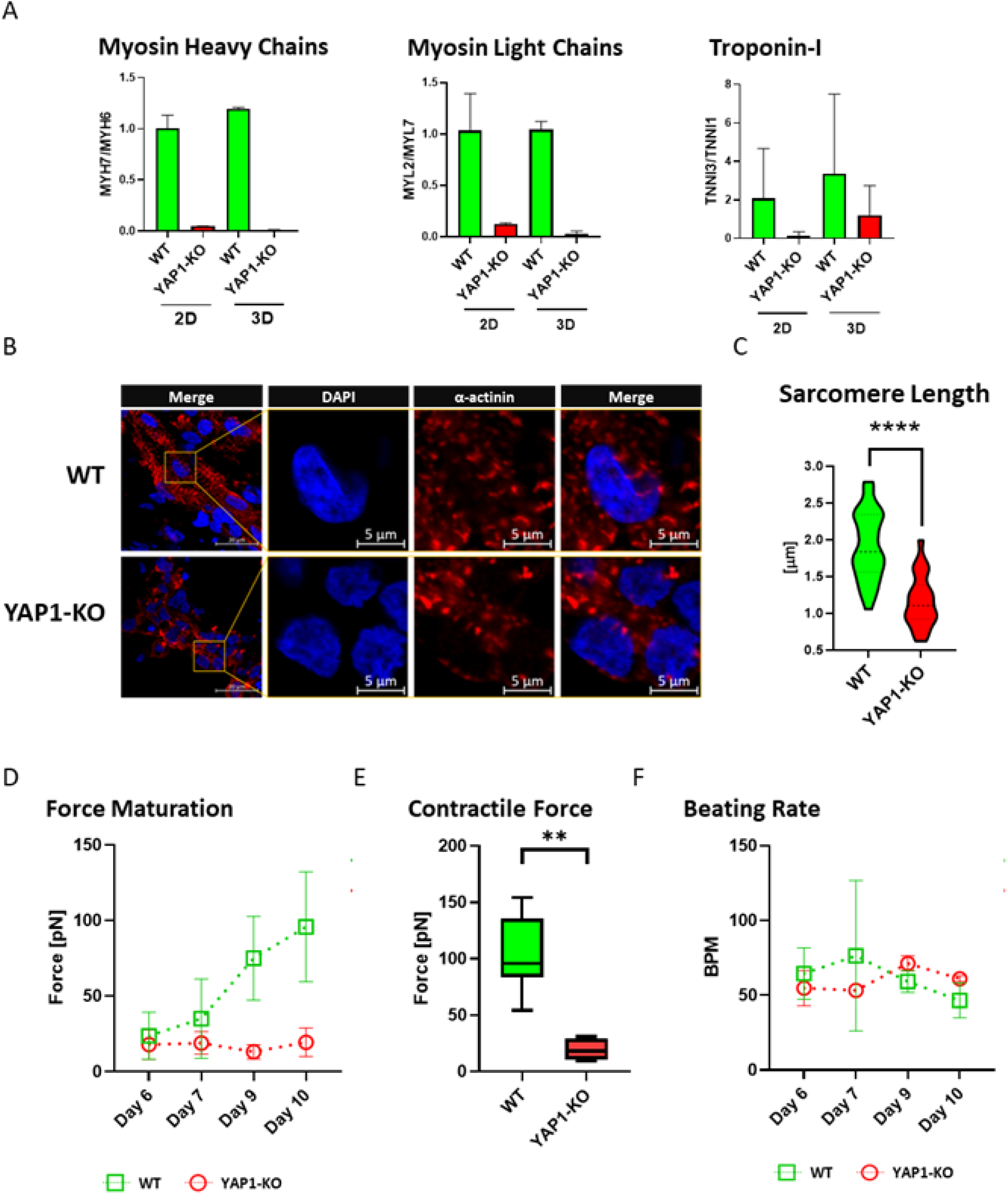
YAP1 promotes contractile force in microtissue model. Cardiomyocytes differentiated from WT and YAP1 deficient induced pluripotent stem cells (iPSCs) for 30 days were cast in 3D microtissues or kept in 2D adherent conditions for further ten days. Maturation associated changes in gene expression visualized as ratios left to right: myosin heavy chains isoforms (MYH7/MYH6), myosin light chains (MYL2/MYL7), and cardiac troponin I isoforms (TNNI3/TNNI1) expression in WT and YAP1-KO hIPSC-CMs at day 40 of differentiation in 2D and 3D cultures was quantified by RT-PCR (n=2) (A). Longitudinal sections of 3D microtissues stained at day 10 post casting for alpha-actinin (red) counterstained with DAPI (blue) (B). Analysis of sarcomere length at day 10 post casting (*P*<0.0001, unpaired *t* test, n=63) (C). Cuore platform (Optics11 Life) was used to measure development of contraction force at different time points up to ten days following casting (D). Contractile force measurements at day 10 post casting (*P*<0.01, unpaired *t* test, n=13) (E). Beating rate measurements up to 10 days post casting (F).

Finally, we live-monitored contractile force up to 10 days post casting into platform. While the force produced by control microtissues increased fourfold over time (from 23.73±15.57 pN at day 6 to 95.81±36.32 pN at day 10), the force of YAP1-KO cardiomyocytes did not change significantly with time in culture (from 17.91±9.863 pN at day 6 to 19.43±9.475 pN at day 10) (Figure 6B, left). The comparison of contractile force between control and YAP1 deficient CMs at day 10 - when the power contraction curve in control cells started to plateau – revealed nearly 5-fold difference in contractile force (Figure 6B, middle). Initially we did not observe big differences in the beating rate between control (64.57±17.15 BPM) and YAP1-KO (54.75±11.73 BPM) on day 6 after EHT formation. Over time the beating rate diverged. While control CMs slowed down slightly (46.60±11.61 BPM at day 10), which is consistent with our observations in cardiac organoids ^58^, the beating rate of YAP1 deficient CMs slightly increased (61±2 BPM) (Figure 6C).

In conclusion, these data indicate that YAP1 is pivotal for maturation of both hESC and hiPSC derived CMs and development of contractile force in 3D EHT constructs.

## Discussion

The enthusiasm of leveraging Hippo pathway in heart regeneration springs from two major discoveries in animal models; (i) the loss of cardiomyocyte proliferation in YAP1 deficient embryonic heart ^16^ and (ii) the expansion of postnatal cardiomyocytes by overexpression of Hippo-insulated constitutively active YAP1 protein (YAP1 S127A) and its variants ^15, 16, 59^. The reactivation of the endogenous protein *in vivo* or the overexpression of unmodified YAP1 *in vitro* does not lead to dramatic cardiomyocyte expansion, but hypertrophy ^20, 21^, which could be relevant for the integration of new cardiomyocytes into muscle tissue. We here used YAP1^-/-^ hESC and hiPSC lines ^14, 31^ as a source of genuine YAP1 deficient cardiomyocytes to explore the function of endogenous YAP1 in the maturation and the adaptive response to pathological conditions.

### Endogenous YAP1 is an important driver of cardiac maturation

Although the protein is strictly necessary for cardio genesis *in vivo* ^18^, the absence of YAP1 posed no barrier to cardiomyocyte differentiation in our *in vitro* experimental model. On the contrary, YAP1 deficient cells differentiated into contractile cardiomyocytes with higher efficiency than WT controls. This result is in line with the need of YAP1 inactivation to mesoderm induction *in vitro* ^36^. On the other hand, the absence of YAP1 hindered the maturation of pluripotent stem cell-derived cardiomyocytes. This result adds to previous reports on the effect of Hippo-mediated constitutively active YAP1 inducing the increased expression of fetal genes ^23^ and to the recent observation that an additional set of chromatin binding sites, which is not accessible to the endogenous protein, might be – instead - accessed by constitutively active YAP1 ^60^.

Increased proliferation is a well-established phenomenon in YAP1 constitutively active gain of function models, but the extent of YAP1 contribution to the basal proliferation rate of cardiomyocytes is less understood. Reduced CM proliferation rate after shRNA-mediated YAP1 silencing in hiPSC-CMs was lately reported ^61^. Our analysis of the proliferation rate in YAP1 deficient cardiomyocytes revealed no significant differences. We attribute this discrepancy to the very high baseline proliferation rate of hiPSC-CMs in Diez-Cuñado study. We and other groups reported nearly one order of magnitude lower baseline proliferation rate of hPSC-CMs ^62–64^. It is thus likely that while the proliferation can be increased by endogenous YAP1 reactivation ^62^ or constitutively active YAP1 expression ^15, 16^, the baseline proliferation rate is YAP1 independent.

Since YAP1 effects are conferred by its interaction with transcriptional partners ^11^, we analyzed genome wide DNA-binding patterns of YAP1 in control cells and compared the transcriptional landscape of normal and YAP1 deficient cardiomyocytes. YAP1 binding to genomic regulatory sequences was associated with increased gene expression confirming the role of endogenous YAP1 as a transcription activator, which is in line with previous published work of constitutively active YAP1 ^60^. Subsets of gene ontologies, which were most repressed by YAP1 absence, govern muscle and sarcomere development. In addition to members of Igf and Wnt family ^16^, previously reported system wide regulators of muscle and sarcomere development in mouse, we observed the repression of thin filament, thick filament, and Z-disc proteins associated with developmental defects and cardiomyopathies (LMOD2, MYH7, MYBPC, CRYAB) ^65–69^.

### Endogenous YAP1 promotes structural and electrophysiological maturation

While some evidence of YAP1 proliferation-independent function on myofibril assembly was described in the fly ^70^ and smaller cardiomyocytes are found in the heart of YAP1 cKO mice ^71^, to our knowledge we bring the first observation of YAP1-regulated myofibril genesis in human *in vitro* cardiomyocytes. In addition, sarcomere maturation was compromised in YAP1 deficient cardiomyocytes but was rescued by YAP1 ectopic re-expression. These observations highlight the difference in function of endogenous and Hippo-insulated constitutively active YAP1. Endogenous YAP1 mediates positive effects on cardiomyocyte maturation induced by metabolic changes ^71^, but the overexpression of Hippo-insulated constitutively active YAP1 either leads to disruption of sarcomere as a prerequisite for cardiomyocyte proliferation ^72^ or does not improve sarcomere maturation ^23, 60^.

Sarcomere development and maturation are interconnected with changes in EC-coupling during heart development. Initially, the combined action of NCX1 and L type Ca^2+^ channel (LTCC, ICaL) start oscillatory Ca^2+^ transients, which later develop into regular oscillations, beat frequency increases and its coordination is delegated to pacemaker cells in sinoatrial node (SAN) using mainly funny current (If) flow through HCN channels to set the frequency ^73–75^. While YAP1 deficiency causes no observable changes prior to E7.5 (start of detectable heart contractions), at day E9.5 heart rate of YAP1 deficient cardiomyocytes slows down before embryonic lethality occurs between days E10.5-E17 ^16^. While our model of YAP1 deficient cardiomyocytes cannot faithfully recapitulate heart development, several aspects seem to be recapitulated. First, genes regulating sodium and calcium currents (ICaL) connected with maturation were repressed in YAP1-KO cardiomyocytes. Next, the spontaneous beating rate of YAP1-KO cardiomyocytes was significantly slower, result well explained by reduced HCN4 gene expression, If density and If open probability. Of note, Hippo-YAP1 axis has been shown to keep homeostasis in sinoatrial node ^76^.

Furthermore, YAP1 deficient cardiomyocytes displayed more immature Ca^2+^ handling properties with reduced SR Ca^2+^ content and slower Ca^2+^ transient decay suggesting decreased efficiency of Ca^2+^ removal systems (mainly SERCA2a and NCX1) in spite of unchanged protein levels of both SERCA2 and NCX1; analogously, the maximal conductance of I_NCX_ was not affected by YAP1 deficiency. Our understanding of the effects of YAP1 transcriptional activity on calcium homeostasis is mixed. While it has been shown that its transcriptional activity through TEAD1 regulates SERCA2a expression and is necessary for adult heart function ^77^, its overexpression leads to heart failure through the repression of Serca2a expression ^22^. In addition, a recent study showed that YAP1 Ca^2+^ transient decay depended on YAP1 activity ^71^. Our study supports the case that the activity of endogenous YAP1 is essential for the maturation of Ca^2+^ handling apparatus.

In postnatal heart both cardiomyocyte proliferation and Hippo regulated YAP1 activity decreases ^18^. Hypertrophic growth then becomes the dominant response of cardiomyocytes to augmented mechanical demands or injury, which is characterized by increased strain and fibronectin deposition *in vivo*. In those conditions, newly induced nuclear YAP1 localization is accompanied by cardiomyocyte hypertrophy ^20, 21^. Cytoskeletal tension was described to regulate YAP1 in Hippo independent pathway through Rho ^9^. Indeed human *in vitro* derived cardiomyocytes recapitulated YAP1 nuclear shuttling and hypertrophic growth in response to both mechanical actuation and fibronectin deposition *in vitro* in a cytoskeleton tension-dependent fashion. YAP1 deficiency hindered this response which is in line with *in vivo* observations from mouse cKO models of myocardial infarction and pressure overload ^20, 21^.

Engineered heart tissues (EHTs) are advanced 3D models suited to assess the functionality of *in vitro* prepared microtissues ^35, 78^. Using EHTs generated from YAP1 deficient hiPSCs, we observed the dramatic decrease in the ability to produce force due to YAP1 genetic depletion. Moreover, we confirmed both the decreased quantity of myofibrils and delayed sarcomere maturation in the same 3D microtissues. To note, the spontaneous beating rate was not significantly altered in YAP1-KO EHTs in comparison to WT; the apparent discrepancy with single cell measurements highlights potential compensatory mechanisms in the 3D model to overcome the huge reduction of the pacemaker current under YAP1- deficiency.

Taken together, our data obtained in both hESCs and hiPSCs in which YAP1 expression has been genetically abolished, demonstrate that YAP1 contributes to cardiomyocyte maturation. Its absence disrupts myofibril alignment and reduces their number. Our results reveal a novel role for Hippo downstream effector in the maturation of cardiomyocyte contractile apparatus and electrophysiological properties. The net effect is the inability of YAP1 deficient cardiomyocytes to respond to physiological stimuli by compensatory growth and reduced force development.

Based on the presented data, we speculate that YAP1 exhibits different modes of action based on the type of activation. Sarcomere disassembly and cardiomyocyte proliferation are effects of Hippo-insulated constitutively active YAP1 S127A. Maturation of sarcomere, electrophysiology, and hypertrophic growth are consequences of the activation of WT version of YAP1. Phased activation of these distinct subsets could lead to two step therapy where Hippo-insulated YAP1 proliferative activity is followed by endogenous-like activated maturation.

## Acknowledgments

Prof. Andrea Barbuti’s contribution was essential in starting electrophysiological measurements of WT and YAP1-deficient cardiomyocytes. As he was sadly not able to see the final results, we would like to acknowledge and remember him for his enthusiasm, expertise, and genuinely good human nature, which inspired us; we will carry it to the future. Further, the authors wish to express their gratitude to Miguel Ramalho-Santos and Han Qin for kindly providing the human embryonic stem cell lines, Prof. Giorgio Giurato from University of Salerno, Italy for Bioinformatics analysis, CELLIM imaging facility for access to SIM microscopy, and Romana Vlckova, Helena Durikova, Helena Skalova, and Hana Dulova from FNUSA-ICRC for the technical support.

## References

1. Sadek H, Olson EN. Toward the Goal of Human Heart Regeneration. Cell Stem Cell. 2020;26:7–16.

2. Garbern JC, Lee RT. Heart regeneration: 20 years of progress and renewed optimism. Dev Cell. 2022;57:424–439.

3. Kawaguchi S, Soma Y, Nakajima K, Kanazawa H, Tohyama S, Tabei R, Hirano A, Handa N, Yamada Y, Okuda S, Hishikawa S, Teratani T, Kunita S, Kishino Y, Okada M, Tanosaki S, Someya S, Morita Y, Tani H, Kawai Y, Yamazaki M, Ito A, Shibata R, Murohara T, Tabata Y, Kobayashi E, Shimizu H, Fukuda K, Fujita J. Intramyocardial Transplantation of Human iPS Cell-Derived Cardiac Spheroids Improves Cardiac Function in Heart Failure Animals. JACC Basic Transl Sci. 2021;6:239–254.

4. Wang J, Liu S, Heallen T, Martin JF. The Hippo pathway in the heart: pivotal roles in development, disease, and regeneration. Nat Rev Cardiol. 2018;15:672–684.

5. Zanconato F, Forcato M, Battilana G, Azzolin L, Quaranta E, Bodega B, Rosato A, Bicciato S, Cordenonsi M, Piccolo S. Genome-wide association between YAP/TAZ/TEAD and AP-1 at enhancers drives oncogenic growth. Nat Cell Biol. 2015;17:1218–27.

6. Yui S, Azzolin L, Maimets M, Pedersen MT, Fordham RP, Hansen SL, Larsen HL, Guiu J, Alves MRP, Rundsten CF, Johansen JV, Li Y, Madsen CD, Nakamura T, Watanabe M, Nielsen OH, Schweiger PJ, Piccolo S, Jensen KB. YAP/TAZ-Dependent Reprogramming of Colonic Epithelium Links ECM Remodeling to Tissue Regeneration. Cell Stem Cell. 2018;22:35–49.e7.

7. Camargo FD, Gokhale S, Johnnidis JB, Fu D, Bell GW, Jaenisch R, Brummelkamp TR. YAP1 increases organ size and expands undifferentiated progenitor cells. Curr Biol CB. 2007;17:2054–60.

8. Nardone G, Oliver-De La Cruz J, Vrbsky J, Martini C, Pribyl J, Skládal P, Pešl M, Caluori G, Pagliari S, Martino F, Maceckova Z, Hajduch M, Sanz-Garcia A, Pugno NM, Stokin GB, Forte G. YAP regulates cell mechanics by controlling focal adhesion assembly. Nat Commun. 2017;8:15321–15321.

9. Dupont S, Morsut L, Aragona M, Enzo E, Giulitti S, Cordenonsi M, Zanconato F, Le Digabel J, Forcato M, Bicciato S, Elvassore N, Piccolo S. Role of YAP/TAZ in mechanotransduction. Nature. 2011;474:179–83.

10. Ren F, Zhang L, Jiang J. Hippo signaling regulates Yorkie nuclear localization and activity through 14-3-3 dependent and independent mechanisms. Dev Biol. 2010;337:303–312.

11. Zhao B, Ye X, Yu J, Li L, Li W, Li S, Yu J, Lin JD, Wang C-Y, Chinnaiyan AM, Lai Z-C, Guan K-L. TEAD mediates YAP-dependent gene induction and growth control. Genes Dev. 2008;22:1962–1971.

12. Cho YS, Jiang J. Hippo-Independent Regulation of Yki/Yap/Taz: A Non-canonical View. Front Cell Dev Biol. 2021;9:658481.

13. Feng X, Degese MS, Iglesias-Bartolome R, Vaque JP, Molinolo AA, Rodrigues M, Zaidi MR, Ksander BR, Merlino G, Sodhi A, Chen Q, Gutkind JS. Hippo-Independent Activation of YAP by the GNAQ Uveal Melanoma Oncogene through a Trio-Regulated Rho GTPase Signaling Circuitry. Cancer Cell. 2014;25:831–845.

14. Qin H, Hejna M, Liu Y, Percharde M, Wossidlo M, Blouin L, Durruthy-Durruthy J, Wong P, Qi Z, Yu J, Qi LS, Sebastiano V, Song JS, Ramalho-Santos M. YAP Induces Human Naive Pluripotency. Cell Rep. 2016;14:2301–12.

15. von Gise A, Lin Z, Schlegelmilch K, Honor LB, Pan GM, Buck JN, Ma Q, Ishiwata T, Zhou B, Camargo FD, Pu WT. YAP1, the nuclear target of Hippo signaling, stimulates heart growth through cardiomyocyte proliferation but not hypertrophy. Proc Natl Acad Sci U S A. 2012;109:2394–9.

16. Xin M, Kim Y, Sutherland LB, Qi X, McAnally J, Schwartz RJ, Richardson JA, Bassel-Duby R, Olson EN. Regulation of Insulin-Like Growth Factor Signaling by Yap Governs Cardiomyocyte Proliferation and Embryonic Heart Size. Sci Signal. 2011;4:ra70.

17. Chen Z, Friedrich GA, Soriano P. Transcriptional enhancer factor 1 disruption by a retroviral gene trap leads to heart defects and embryonic lethality in mice. Genes Dev. 1994;8:2293–2301.

18. Heallen T, Zhang M, Wang J, Bonilla-Claudio M, Klysik E, Johnson RL, Martin JF. Hippo Pathway Inhibits Wnt Signaling to Restrain Cardiomyocyte Proliferation and Heart Size. Science. 2011;332:458–461.

19. Liu R, Jagannathan R, Li F, Lee J, Balasubramanyam N, Kim BS, Yang P, Yechoor VK, Moulik M. Tead1 is required for perinatal cardiomyocyte proliferation. PLoS ONE. 2019;14:e0212017.

20. Del Re DP, Yang Y, Nakano N, Cho J, Zhai P, Yamamoto T, Zhang N, Yabuta N, Nojima H, Pan D, Sadoshima J. Yes-associated Protein Isoform 1 (Yap1) Promotes Cardiomyocyte Survival and Growth to Protect against Myocardial Ischemic Injury. J Biol Chem. 2013;288:3977–3988.

21. Byun J, Re DPD, Zhai P, Ikeda S, Shirakabe A, Mizushima W, Miyamoto S, Brown JH, Sadoshima J. Yes-associated protein (YAP) mediates adaptive cardiac hypertrophy in response to pressure overload. J Biol Chem. 2019;294:3603–3617.

22. Tsika RW, Ma L, Kehat I, Schramm C, Simmer G, Morgan B, Fine DM, Hanft LM, McDonald KS, Molkentin JD, Krenz M, Yang S, Ji J. TEAD-1 Overexpression in the Mouse Heart Promotes an Age-dependent Heart Dysfunction. J Biol Chem. 2010;285:13721– 13735.

23. Ikeda S, Mizushima W, Sciarretta S, Abdellatif M, Zhai P, Mukai R, Fefelova N, Oka S, Nakamura M, Del Re DP, Farrance I, Park JY, Tian B, Xie L-H, Kumar M, Hsu C-P, Sadayappan S, Shimokawa H, Lim D-S, Sadoshima J. Hippo Deficiency Leads to Cardiac Dysfunction Accompanied by Cardiomyocyte De-Differentiation During Pressure Overload. Circ Res. 2019;124:292–305.

24. Gabisonia K, Prosdocimo G, Aquaro GD, Carlucci L, Zentilin L, Secco I, Ali H, Braga L, Gorgodze N, Bernini F, Burchielli S, Collesi C, Zandonà L, Sinagra G, Piacenti M, Zacchigna S, Bussani R, Recchia FA, Giacca M. MicroRNA therapy stimulates uncontrolled cardiac repair after myocardial infarction in pigs. Nature. 2019;569:418– 422.

25. Torrini C, Cubero RJ, Dirkx E, Braga L, Ali H, Prosdocimo G, Gutierrez MI, Collesi C, Licastro D, Zentilin L, Mano M, Zacchigna S, Vendruscolo M, Marsili M, Samal A, Giacca M. Common Regulatory Pathways Mediate Activity of MicroRNAs Inducing Cardiomyocyte Proliferation. Cell Rep. 2019;27:2759–2771.e5.

26. Papait R, Cattaneo P, Kunderfranco P, Greco C, Carullo P, Guffanti A, Viganò V, Stirparo GG, Latronico MVG, Hasenfuss G, Chen J, Condorelli G. Genome-wide analysis of histone marks identifying an epigenetic signature of promoters and enhancers underlying cardiac hypertrophy. Proc Natl Acad Sci. 2013;110:20164–20169.

27. Yang Y, Del Re DP, Nakano N, Sciarretta S, Zhai P, Park J, Sayed D, Shirakabe A, Matsushima S, Park Y, Tian B, Abdellatif M, Sadoshima J. miR-206 Mediates YAP- Induced Cardiac Hypertrophy and Survival. Circ Res. 2015;117:891–904.

28. Perestrelo AR, Silva AC, Oliver-De La Cruz J, Martino F, Horváth V, Caluori G, Polanský O, Vinarský V, Azzato G, de Marco G, Žampachová V, Skládal P, Pagliari S, Rainer A, Pinto-do-Ó P, Caravella A, Koci K, Nascimento DS, Forte G. Multiscale Analysis of Extracellular Matrix Remodeling in the Failing Heart. Circ Res. 2021;128:24–38.

29. Vrbský J, Vinarský V, Perestrelo AR, De La Cruz JO, Martino F, Pompeiano A, Izzi V, Hlinomaz O, Rotrekl V, Sudol M, Pagliari S, Forte G. Evidence for discrete modes of YAP1 signaling via mRNA splice isoforms in development and diseases. Genomics. 2021;113:1349–1365.

30. Martino F, Varadarajan NM, Perestrelo AR, Hejret V, Durikova H, Vukic D, Horvath V, Cavalieri F, Caruso F, Albihlal WS, Gerber AP, O’Connell MA, Vanacova S, Pagliari S, Forte G. The mechanical regulation of RNA binding protein hnRNPC in the failing heart. Sci Transl Med. 2022;14:eabo5715.

31. Zeevaert K, Goetzke R, Elsafi Mabrouk MH, Schmidt M, Maaßen C, Henneke A-C, He C, Gillner A, Zenke M, Wagner W. YAP1 is essential for self-organized differentiation of pluripotent stem cells. Biomater Adv. 2023;146:213308.

32. Lian X, Hsiao C, Wilson G, Zhu K, Hazeltine LB, Azarin SM, Raval KK, Zhang J, Kamp TJ, Palecek SP. Robust cardiomyocyte differentiation from human pluripotent stem cells via temporal modulation of canonical Wnt signaling. Proc Natl Acad Sci U S A. 2012;109:E1848–E1857.

33. Palchesko RN, Zhang L, Sun Y, Feinberg AW. Development of Polydimethylsiloxane Substrates with Tunable Elastic Modulus to Study Cell Mechanobiology in Muscle and Nerve. PLOS ONE. 2012;7:e51499.

34. Vinarský V, Martino F, Forte G, Šleichrt J, Rada V, Kytýř D. DEFORMATION RESPONSE OF POLYDIMETHYLSILOXANE SUBSTRATES SUBJECTED TO UNIAXIAL QUASI-STATIC LOADING. Acta Polytech CTU Proc. 2019;25:79–82.

35. Iuliano A, Haalstra M, Raghuraman R, Bielawski K, Bholasing AP, van der Wal E, de Greef JC, Pijnappel WWMP. Real-time and Multichannel Measurement of Contractility of hiPSC-Derived 3D Skeletal Muscle using Fiber Optics-Based Sensing. Adv Mater Technol. 2023;8:2300845.

36. Pagliari S, Vinarsky V, Martino F, Perestrelo AR, Oliver De La Cruz J, Caluori G, Vrbsky J, Mozetic P, Pompeiano A, Zancla A, Ranjani SG, Skladal P, Kytyr D, Zdráhal Z, Grassi G, Sampaolesi M, Rainer A, Forte G. YAP–TEAD1 control of cytoskeleton dynamics and intracellular tension guides human pluripotent stem cell mesoderm specification. Cell Death Differ. 2021;28:1193–1207.

37. Vinarsky V, Krivanek J, Rankel L, Nahacka Z, Barta T, Jaros J, Andera L, Hampl A. Human embryonic and induced pluripotent stem cells express TRAIL receptors and can be sensitized to TRAIL-induced apoptosis. Stem Cells Dev. 2013;

38. Berg S, Kutra D, Kroeger T, Straehle CN, Kausler BX, Haubold C, Schiegg M, Ales J, Beier T, Rudy M, Eren K, Cervantes JI, Xu B, Beuttenmueller F, Wolny A, Zhang C, Koethe U, Hamprecht FA, Kreshuk A. ilastik: interactive machine learning for (bio)image analysis. Nat Methods. 2019;16:1226–1232.

39. Stirling DR, Swain-Bowden MJ, Lucas AM, Carpenter AE, Cimini BA, Goodman A. CellProfiler 4: improvements in speed, utility and usability. BMC Bioinformatics. 2021;22:433.

40. Schindelin J, Arganda-Carreras I, Frise E, Kaynig V, Longair M, Pietzsch T, Preibisch S, Rueden C, Saalfeld S, Schmid B, Tinevez J-Y, White DJ, Hartenstein V, Eliceiri K, Tomancak P, Cardona A. Fiji: an open-source platform for biological-image analysis. Nat Methods. 2012;9:676–682.

41. Langmead B, Trapnell C, Pop M, Salzberg SL. Ultrafast and memory-efficient alignment of short DNA sequences to the human genome. Genome Biol. 2009;10:R25–R25.

42. Uusküla-Reimand L, Hou H, Samavarchi-Tehrani P, Rudan MV, Liang M, Medina-Rivera A, Mohammed H, Schmidt D, Schwalie P, Young EJ, Reimand J, Hadjur S, Gingras A-C, Wilson MD. Topoisomerase II beta interacts with cohesin and CTCF at topological domain borders. Genome Biol. 2016;17:182–182.

43. Heinz S, Benner C, Spann N, Bertolino E, Lin YC, Laslo P, Cheng JX, Murre C, Singh H, Glass CK. Simple Combinations of Lineage-Determining Transcription Factors Prime cis-Regulatory Elements Required for Macrophage and B Cell Identities. Mol Cell. 2010;38:576–589.

44. Chen T, Dent SYR. Chromatin modifiers and remodellers: regulators of cellular differentiation. Nat Rev Genet. 2014;15:93–106.

45. Zambelli F, Pesole G, Pavesi G. PscanChIP: finding over-represented transcription factor-binding site motifs and their correlations in sequences from ChIP-Seq experiments. Nucleic Acids Res. 2013;41:W535–W543.

46. Kim D, Pertea G, Trapnell C, Pimentel H, Kelley R, Salzberg SL. TopHat2: accurate alignment of transcriptomes in the presence of insertions, deletions and gene fusions. Genome Biol. 2013;14:R36.

47. Anders S, Pyl PT, Huber W. HTSeq—a Python framework to work with high- throughput sequencing data. Bioinformatics. 2015;31:166–169.

48. Benzoni P, Arici M, Giannetti F, Cospito A, Prevostini R, Volani C, Fassina L, Rosato-Siri MD, Metallo A, Gennaccaro L, Suffredini S, Foco L, Mazzetti S, Calogero A, Cappelletti G, Leibbrandt A, Elling U, Broso F, Penninger JM, Pramstaller PP, Piubelli C, Bucchi A, Baruscotti M, Rossini A, Rocchetti M, Barbuti A. Striatin knock out induces a gain of function of INa and impaired Ca2+ handling in mESC-derived cardiomyocytes. Acta Physiol Oxf Engl. 2024;e14160.

49. Fishilevich S, Nudel R, Rappaport N, Hadar R, Plaschkes I, Iny Stein T, Rosen N, Kohn A, Twik M, Safran M, Lancet D, Cohen D. GeneHancer: genome-wide integration of enhancers and target genes in GeneCards. Database J Biol Databases Curation. 2017;2017:bax028.

50. Estarás C, Hsu H-T, Huang L, Jones KA. YAP repression of the WNT3 gene controls hESC differentiation along the cardiac mesoderm lineage. Genes Dev. 2017;31:2250– 2263.

51. Wheelwright M, Win Z, Mikkila JL, Amen KY, Alford PW, Metzger JM. Investigation of human iPSC-derived cardiac myocyte functional maturation by single cell traction force microscopy. PLOS ONE. 2018;13:e0194909.

52. Ronaldson-Bouchard K, Ma SP, Yeager K, Chen T, Song L, Sirabella D, Morikawa K, Teles D, Yazawa M, Vunjak-Novakovic G. Advanced maturation of human cardiac tissue grown from pluripotent stem cells. Nature. 2018;556:239–243.

53. Lundy SD, Zhu W-Z, Regnier M, Laflamme MA. Structural and Functional Maturation of Cardiomyocytes Derived from Human Pluripotent Stem Cells. Stem Cells Dev. 2013;22:1991–2002.

54. Frangogiannis NG. Cardiac fibrosis. Cardiovasc Res. 2021;117:1450–1488.

55. Torres WM, Jacobs J, Doviak H, Barlow SC, Zile MR, Shazly T, Spinale FG. Regional and temporal changes in left ventricular strain and stiffness in a porcine model of myocardial infarction. Am J Physiol - Heart Circ Physiol. 2018;315:H958–H967.

56. van Duijvenboden K, de Bakker DEM, Man JCK, Janssen R, Günthel M, Hill MC, Hooijkaas IB, van der Made I, van der Kraak PH, Vink A, Creemers EE, Martin JF, Barnett P, Bakkers J, Christoffels VM. Conserved NPPB+ Border Zone Switches From MEF2- to AP-1-Driven Gene Program. Circulation. 2019;140:864–879.

57. Guo Y, Pu WT. Cardiomyocyte Maturation: New Phase in Development. Circ Res. 2020;126:1086–1106.

58. Ergir E, Oliver-De La Cruz J, Fernandes S, Cassani M, Niro F, Pereira-Sousa D, Vrbský J, Vinarský V, Perestrelo AR, Debellis D, Vadovičová N, Uldrijan S, Cavalieri F, Pagliari S, Redl H, Ertl P, Forte G. Generation and maturation of human iPSC-derived 3D organotypic cardiac microtissues in long-term culture. Sci Rep. 2022;12:17409.

59. Lin Z, Gise A von, Zhou P, Gu F, Ma Q, Jiang J, Yau AL, Buck JN, Gouin KA, Gorp PRR van, Zhou B, Chen J, Seidman JG, Wang D, Pu WT. Cardiac-Specific YAP Activation Improves Cardiac Function and Survival in an Experimental Murine MI Model. Circ Res. 2014;115:354.

60. Monroe TO, Hill MC, Morikawa Y, Leach JP, Heallen T, Cao S, Krijger PHL, de Laat W, Wehrens XHT, Rodney GG, Martin JF. YAP Partially Reprograms Chromatin Accessibility to Directly Induce Adult Cardiogenesis In Vivo. Dev Cell. 2019;48:765–779.e7.

61. Diez-Cuñado M, Wei K, Bushway PJ, Maurya MR, Perera R, Subramaniam S, Ruiz- Lozano P, Mercola M. miRNAs that Induce Human Cardiomyocyte Proliferation Converge on the Hippo Pathway. Cell Rep. 2018;23:2168–2174.

62. Neininger AC, Dai X, Liu Q, Burnette DT. The Hippo pathway regulates density- dependent proliferation of iPSC-derived cardiac myocytes. Sci Rep. 2021;11:17759.

63. Buikema JW, Lee S, Goodyer WR, Maas RG, Chirikian O, Li G, Miao Y, Paige SL, Lee D, Wu H, Paik DT, Rhee S, Tian L, Galdos FX, Puluca N, Beyersdorf B, Hu J, Beck A, Venkamatran S, Swami S, Wijnker P, Schuldt M, Dorsch LM, van Mil A, Red-Horse K, Wu JY, Geisen C, Hesse M, Serpooshan V, Jovinge S, Fleischmann BK, Doevendans PA, van der Velden J, Garcia KC, Wu JC, Sluijter JPG, Wu SM. Wnt Activation and Reduced Cell-Cell Contact Synergistically Induce Massive Expansion of Functional Human iPSC- Derived Cardiomyocytes. Cell Stem Cell. 2020;27:50–63.e5.

64. Park S, Choe M, Yeo H, Han H, Kim J, Chang W, Yun S, Lee H, Lee M. Yes-associated protein mediates human embryonic stem cell-derived cardiomyocyte proliferation: Involvement of epidermal growth factor receptor signaling. J Cell Physiol. 2018;233:7016–7025.

65. Vasilescu C, Colpan M, Ojala TH, Manninen T, Mutka A, Ylänen K, Rahkonen O, Poutanen T, Martelius L, Kumari R, Hinterding H, Brilhante V, Ojanen S, Lappalainen P, Koskenvuo J, Carroll CJ, Fowler VM, Gregorio CC, Suomalainen A. Recessive TMOD1 mutation causes childhood cardiomyopathy. Commun Biol. 2024;7:7.

66. Lay E, Azamian MS, Denfield SW, Dreyer W, Spinner JA, Kearney D, Zhang L, Worley KC, Bi W, Lalani SR. LMOD2-related dilated cardiomyopathy presenting in late infancy. Am J Med Genet A. 2022;188:1858–1862.

67. Watkins H, Conner D, Thierfelder L, Jarcho JA, MacRae C, McKenna WJ, Maron BJ, Seidman JG, Seidman CE. Mutations in the cardiac myosin binding protein-C gene on chromosome 11 cause familial hypertrophic cardiomyopathy. Nat Genet. 1995;11:434–437.

68. Maloyan A, Sanbe A, Osinska H, Westfall M, Robinson D, Imahashi K, Murphy E, Robbins J. Mitochondrial dysfunction and apoptosis underlie the pathogenic process in alpha-B-crystallin desmin-related cardiomyopathy. Circulation. 2005;112:3451–3461.

69. Fananapazir L, Dalakas MC, Cyran F, Cohn G, Epstein ND. Missense mutations in the beta-myosin heavy-chain gene cause central core disease in hypertrophic cardiomyopathy. Proc Natl Acad Sci U S A. 1993;90:3993–3997.

70. Kaya-Çopur A, Marchiano F, Hein MY, Alpern D, Russeil J, Luis NM, Mann M, Deplancke B, Habermann BH, Schnorrer F. The Hippo pathway controls myofibril assembly and muscle fiber growth by regulating sarcomeric gene expression. eLife. 2021;10:e63726.

71. Murphy SA, Miyamoto M, Kervadec A, Kannan S, Tampakakis E, Kambhampati S, Lin BL, Paek S, Andersen P, Lee D-I, Zhu R, An SS, Kass DA, Uosaki H, Colas AR, Kwon C. PGC1/PPAR drive cardiomyocyte maturation at single cell level via YAP1 and SF3B2. Nat Commun. 2021;12:1648.

72. Ahuja P, Perriard E, Perriard J-C, Ehler E. Sequential myofibrillar breakdown accompanies mitotic division of mammalian cardiomyocytes. J Cell Sci. 2004;117:3295–3306.

73. Tyser RC, Miranda AM, Chen C, Davidson SM, Srinivas S, Riley PR. Calcium handling precedes cardiac differentiation to initiate the first heartbeat. eLife. 2016;5:e17113.

74. Jia BZ, Qi Y, Wong-Campos JD, Megason SG, Cohen AE. A bioelectrical phase transition patterns the first vertebrate heartbeats. Nature. 2023;622:149–155.

75. Ji RP, Phoon CKL, Aristizábal O, McGrath KE, Palis J, Turnbull DH. Onset of Cardiac Function During Early Mouse Embryogenesis Coincides With Entry of Primitive Erythroblasts Into the Embryo Proper. Circ Res. 2003;92:133–135.

76. Zheng M, Li RG, Song J, Zhao X, Tang L, Erhardt S, Chen W, Nguyen BH, Li X, Li M, Wang J, Evans SM, Christoffels VM, Li N, Wang J. Hippo-Yap signaling maintains sinoatrial node homeostasis. Circulation. 2022;146:1694–1711.

77. Liu R, Lee J, Kim BS, Wang Q, Buxton SK, Balasubramanyam N, Kim JJ, Dong J, Zhang A, Li S, Gupte AA, Hamilton DJ, Martin JF, Rodney GG, Coarfa C, Wehrens XHT, Yechoor VK, Moulik M. Tead1 is required for maintaining adult cardiomyocyte function, and its loss results in lethal dilated cardiomyopathy. JCI Insight. 2017;2:e93343.

78. Eder A, Vollert I, Hansen A, Eschenhagen T. Human engineered heart tissue as a model system for drug testing. Adv Drug Deliv Rev. 2016;96:214–224.

